# The impact of fast radiation on the phylogeny of *Bactrocera* fruit flies

**DOI:** 10.1101/2021.09.07.459237

**Authors:** Federica Valerio, Nicola Zadra, Omar Rota Stabelli, Lino Ometto

**Affiliations:** Department of Biology and Biotechnology, University of Pavia, 27100 Pavia, Italy; Research and Innovation Centre, Fondazione Edmund Mach, 38010 San Michele all’Adige (TN), Italy; Department of Cellular, Computational and Integrative Biology - CIBIO, University of Trento, 38123 Trento, Italy; Center Agriculture Food Environment - C3A, University of Trento, 38010 San Michele all’Adige (TN), Italy

## Abstract

True fruit flies (Tephritidae) include several species that cause extensive damage to agriculture worldwide. Among them, species of the genus *Bactrocera* are widely studied to understand the traits associated to their invasiveness and ecology. Comparative approaches based on a reliable phylogenetic framework are particularly effective, but, to date, molecular phylogenies of *Bactrocera* are still controversial. Here, we employed a comprehensive genomic dataset to infer a robust backbone phylogeny of eleven representative *Bactrocera* species and two outgroups. We further provide the first genome scaled inference of their divergence using calibrated relaxed clock. The results of our analyses support a closer relationship of *B. dorsalis* to *B. latifrons* than to *B. tryoni*, in contrast to all mitochondrial-based phylogenies. By comparing different evolutionary models, we show that this incongruence likely derives from the fast and recent radiation of these species that occurred around 2 million years ago, which may be associated with incomplete lineage sorting and possibly (ongoing) hybridization. These results can serve as basis for future comparative analyses and highlight the utility of using large datasets and efficient phylogenetic approaches to study the evolutionary history of species of economic importance.

---

Tephritidae, commonly known as true fruit flies, are an incredibly diverse group of phytophagous insects. They include more than 5,000 species in over 500 genera (Bickel *et al.*, 2009), making it one of the largest families among Diptera. Several of these species cause extensive damage to agriculture worldwide, since females lay eggs within the fruit flesh (i.e., pericarp) and the larvae rapidly develop in the fruit, eating it and inducing bacterial and fungal decay. The accompanying economic impacts are considerable: for example, the olive fruit fly, *Bactrocera oleae* (Rossi), has been estimated to cause an annual economic loss of approximately $800 million in the Mediterranean basin alone (Daane & Johnson, 2010). Moreover, the economic impact of these species is exacerbated by their ability to invade new areas and establish viable populations.

Most relevant tephritid pests typically belong to five genera: *Anastrepha* Schiner, *Ceratitis* MacLeay, *Rhagoletis* Loew, *Zeugodacus* Hendel and *Bactrocera* Macquart. The genus *Bactrocera* is the most economically important and comprises over 500 species (Drew, 1989), at least 50 of which are considered important pests (White & Elson-Harris, 1992). Identifying the key ecological and biological traits relevant for their invasiveness and host preference is extremely important to drive effective control strategies. To this aim, comparative approaches based on a reliable phylogenetic framework are particularly effective, as they allow to trace the evolution of species-specific and shared traits and evaluate whether, and to what extent control measures may be applied in related species (Ometto *et al.*, 2013). The *Bactrocera* genus has a controversial history of taxonomy and classification. The taxonomic status of this group has been revised many times and it was established as a genus only in 1989 (Drew, 1989). The genus is subdivided in ~30 subgenera (Drew, 1972), sometimes recognized as distinct genera, e.g. *Dacus* (Drew & Romig, 2016) and *Zeugodacus* (Drew & Hancock, 1999). To date, molecular phylogenies of *Bactrocera* resulted in different relationships depending on the type (nuclear and/or mitochondrial) and number of markers, and on the number of taxa analyzed (Krosch *et al.*, 2012; Muraji & Nakahara, 2001; Nakahara & Muraji, 2007; San Jose *et al.*, 2018; Smith *et al.*, 2003; Virgilio *et al.*, 2015). For instance, *B. dorsalis* is usually considered to be more closely related to *B. tryoni* than to *B. latifrons* (e.g., (Yong *et al.*, 2016; A. Bin Zhang *et al.*, 2010), but recent studies suggested a possible (i.e. unsupported) closer relationship between *B. dorsalis* and *B. latifrons* (Choo *et al.*, 2019; San Jose *et al.*, 2018). This uncertainty is particularly relevant for pest management, since these three species are important pests and different phylogenetic reconstructions may affect the interpretation of comparative studies.

Here, we employed a phylogenomics approach to infer a robust phylogeny of Tephritidae with a focus on Bactrocera and we further calibrated a relaxed clock to estimate their divergences. Our aim is to provide a phylogenetic and chronological backbone that can serve as basis for comparative analyses. Our dataset encompasses the large distribution and diversity of Bactrocera and since it includes species characterized by distinct diets, which range from polyphagy (e.g., *B. dorsalis*) to monophagy (e.g., *B. oleae*), understanding the evolutionary history of this group may also provide precious information on the mode and tempo of the evolution of their biology and ecology.

## Materials and Methods

### Datasets

In our analysis, we included species for which genomic and/or transcriptomic resources were available: *Afrodacus* (*B. jarvisi* (Tryon)), *Daculus* (*B. oleae* (Gmelin)), *Tetradacus* (*B. minax* (Enderlein)), *Bactrocera* (*B. bryoniae* (Tryon), *B. correcta* (Bezzi), *B. dorsalis* (Hendel), *B. latifrons* (Hendel), *B. musae* (Tryon), *B. tryoni* (Froggatt) and *B. zonata* (Saunders)), and *Zeugodacus cucurbitae* (Coquillett). In particular, we analysed datasets of coding sequences (CDS) and of RNA (transcript) sequences for *Ceratitis capitata* (Papanicolaou *et al.*, 2016), *Z. cucurbitae* (Sim & Geib, 2017), *B. dorsalis* (Geib *et al.*, 2014), *B. latifrons* (NCBI accession: MIMC00000000.1), *B. minax* (Wang *et al.*, 2016) and *B. oleae* (Bayega *et al.*, 2020). For the six remaining *Bactrocera* species (*B. bryoniae*, *B. correcta*, *B. jarvisi*, *B. musae*, *B. tryoni*, and *B. zonata*), we downloaded the available RNA-Seq raw reads (see Supplemental Table 1 for SRA accession numbers) and assembled the corresponding transcriptomes using default parameters with Trinity v. 2.7.0 (Grabherr *et al.*, 2011).

### Orthologous gene set identification and alignment

Orthologs across the ten *Bactrocera* species, *Z. cucurbitae* and the outgroup *C. capitata* were identified using a reciprocal-best-hit approach using pairwise BLASTn searches (Camacho *et al.*, 2009) between the *C. capitata* CDS sequences and each of the other datasets. Putative 1:1 orthologs were first aligned using MAFFT (Katoh & Standley, 2013) and any incomplete codon (based on the *C. capitata* sequence) was removed. We then re-aligned the ortholog sets using the PRANK algorithm (Löytynoja & Goldman, 2008) implemented in the tool TranslatorX (Abascal *et al.*, 2010). We minimized bias in our datasets by 1) removing alignments containing sequences with internal stop codons and 2) using a custom perl script to remove problematic and ambiguous alignment regions (Ramasamy *et al.*, 2016). Using this pipeline, we ultimately identified 110 orthologous gene sets across all the twelve species.

We concatenated orthologs to generate an alignment of 189,891 nucleotides (nt) and then translated it using the standard genetic code to obtain an alignment of 63,297 amino acids (aa). We further generated an alignment of 24,885 nt containing only 4-fold degenerate sites.

### Phylogenetic analyses

We inferred phylogenetic relationships using both a maximum likelihood (ML) and a Bayesian approach. To explore possible sources of systematic errors we employed homogenous, heterogeneous, and coalescent aware models of evolution.

We run ML analyses on both the concatenated aa alignment using the PROTGAMMAGTR model, and on the nt alignment partitioned in first, second and third (1+2+3) position, using the GAMMAGTR model implemented in RAxML (Stamatakis, 2014). We also run a ML analysis based on the 4-fold degenerate sites alignment using the GAMMAGTR model. In all cases, node support was calculated by the rapid bootstrap feature of RAxML (100 replicates). We also estimated bootstrap supports using the coalescent-aware analysis of ASTRAL (C. Zhang *et al.*, 2018), which was based on all single ML gene trees obtained by RAxML using the same models of the concatenated analyses for either the nt or the aa sequences. Bootstrap values were estimated by performing either 100 multi-locus bootstrap replicates or gene+site resampling (using the −g option).

The same three datasets, aa, nt (1+2+3), and 4-fold degenerate sites, were used to run Bayesian analyses in Beast v.2.5.1 (Bouckaert *et al.*, 2019). The aminoacidic dataset was analysed with a LG+G4 substitution model. The 4-fold degenerate sites dataset and the codon dataset were analysed with a GTR+G4 substitution model. The codon dataset was split into three partitions, corresponding to the codon positions, setting linked trees across them. We performed additional Bayesian analysis on the aminoacidic dataset using a CAT+GTR model and on the 4-fold degenerate sites dataset using the among-site heterogeneous CAT model with gamma distribution with PhyloBayes (Lartillot & Philippe, 2004). In both cases we run two independent MCMC chains and checked for convergence using the associated tracecomp and bpcomp commands. For the aminoacidic dataset we let both chains run until parameters were stabilized with *maxdiff* =0. For the 4-fold degenerate sites dataset both chains run until parameters were stabilized with *maxdiff* =0, *reldiff* < 0.25 (except for stat, with *reldiff* < 0.4) and the effective sample size was *effsize* > 170 (except for nmode, statalpha, kappa and allocent, with *effsize* between 17 and 85). The summary statistics reached stability at the end of the runs and were periodically visualized with the script graphphylo (https://github.com/wrf/graphphylo).

Bayesian analyses were also run using StarBeast2 (Ogilvie *et al.*, 2017), which employs a multispecies coalescent method to estimate species trees from multiple sequence alignments, as implemented in Beast package and according to the tutorial provided by Taming the Beast (Barido-Sottani *et al.*, 2018). For this analysis we used either the nucleotide or the amino acidic alignments, linking the site models across the gene sets.

BEAST and StarBeast2 analyses had chains run for 10^8^ generations (or until convergence for at least 4×10^7^ generations), sampling trees and parameters every 1,000 generations and inspecting convergence and likelihood plateauing in Tracer. Posterior consensus trees were generated after discarding the first 10% of generations as burn-in. For the StarBeast analysis, single gene trees were loaded and visualized by DensiTree.

### Dating analysis

Because of the numerous incongruences between the species tree obtained by the multi-locus analyses and the single gene trees (see Results section), which could bias the dating analysis, we produced a conservative dataset by: i) limiting the species sample to 10 representative species (*C. capitata*, *Z. cucurbitae, B. bryoniae*, *B. dorsalis*, *B. jarvisi*, *B. latifrons*, *B. minax*, *B. musae*, *B. oleae*, *B. tryoni*) and ii) considering only those 37 genes that produced a ML tree supporting the consensus species tree with minimum ML bootstrap values of 50 at each node.

Divergence times were then estimated by Beast v. 2.5.1 using the 4-fold degenerate sites of the concatenated dataset (11,768 nt). This dataset allowed us to use an instantaneous (neutral) mutation rate as prior. Since mutation rate in Tephritidae is not known yet, we assumed it to be similar to that of *Drosophila* (another Diptera) and used the estimate of 0.0346 (SD = 0.0028) substitutions per base pair per million years provided by (Obbard *et al.*, 2012). Because in *Bactrocera* we assumed eight generations per year (in nature they range from 3-5 of *B. oleae* and sub-tropical *B. dorsalis* populations, to >12 for the tropical species; (Li *et al.*, 2019; Stephens *et al.*, 2007; Theron *et al.*, 2017; Vargas *et al.*, 1997)) and to account for uncertainty, we finally set as prior a normally distributed mean of 0.028 (SD = 0.03). In a second approach, we set a mutation rate lognormal distributed with ‘mean in real space’ M = 0.028 and S = 0.82 (to produce the same 95% quantile – 0.077 – as the normal distribution). For both approaches we performed a model selection to choose the most fitting clock and demographic prior based on the marginal likelihood values with the nested sampling approach implemented in the NS package (Russel *et al.*, 2019). We tested the strict and the LOGN relaxed clock and the Yule and the Birth-Death models, for a total of eight different combinations. Following the recommendations provided by the dedicated Taming the Beast tutorial (Barido-Sottani *et al.*, 2018), sub-chain length was set at 50,000, which corresponds to the length of the MCMC run (i.e., 5×10^7^) divided by the smallest ESS value observed across the eight model runs (i.e., ~1,000), and the number of particles was set at 10. A model was considered favoured over another model if the difference between the two marginal likelihoods (i.e., the Bayes Factor (BF) in log space) was more than twice the sum of the corresponding standard deviations (SD). We run eight different analyses that used different combinations of priors and model settings and performed a model selection to identify the most appropriate for our data. In particular, the nested sampling approach allowed us to estimate the marginal likelihoods of the different models and make pairwise comparisons using the associated Bayes Factor. All models had marginal likelihoods with a standard deviation ranging from 2.5 to 2.9, which was small enough to assess whether a model was favored over another one. In all cases, we employed a GTR+G replacement model and a root prior uniformly distributed between 6 and 65 million years ago (Mya), which correspond to the age of a *Ceratitis* fly fossil (Norrbom, 1994) and of the Schizophora radiation (Junqueira *et al.*, 2016; Wiegmann *et al.*, 2011). Because the Bayesian phylogenetic analysis on the concatenated 4-fold degenerate sites resulted in a topology incongruent to the one supported by all other ML and Bayesian analyses (see Results), the species tree was fixed according to the latter consensus topology.

All analyses were run twice, with chains run for 5×10^7^ generations, sampling trees and parameters every 1,000 generations and inspecting convergence and likelihood plateauing in Tracer. Both chains resulted well mixed, with average effective sample size (ESS) values across posterior values being well above 200. The consensus trees (Maximum Clade Credibility trees) were generated after discarding the first 20% of generations as burn-in.

## Results and Discussion

### Phylogenetic analyses a closer affinity of *B. dorsalis* to *B. latifrons* than to *B. tryoni*

The results of our analyses strongly indicate that *B. dorsalis* is more closely related to *B. latifrons* than to *B. tryoni* (Fig. 1). This relationship is highly supported according to both the ML bootstrap values (≥ 98, Fig. 1A and Supplementary Figure 1) and the Bayesian posterior probabilities obtained using both BEAST and PhyloBayes analyses (PP = 1, Fig. 1A and Supplementary Figures 2-4), no matter whether based on the codon or aminoacidic alignments. The only support for the relationship ((*B. dorsalis*, *B. tryoni*), *B. latifrons*) comes from the Bayesian analysis run in Beast2 and based on the concatenated alignment of the 4-fold degenerate sites (Supplementary Figure 5). This latter dataset contains sites under (nearly) neutral evolution and therefore is suitable for divergence time estimates because it allows a calibration using instantaneous mutation rates; it is however more prone to saturation compared with the nucleotide and amino acidic datasets. When the 4-fold degenerate sites dataset was analyzed in PhyloBayes (which is also a Bayesian implementation) using the among site heterogeneous CAT model instead of an among-site homogenous model as in Beast2, we obtain the same topology obtained by the aforementioned Bayesian and ML analyses on the nucleotide and amino acidic datasets (Supplementary Figure 6; see also Supplementary Figure 7 for the results of the ML analysis on the same dataset). Heterogeneous models such as CAT can accommodate for systematic errors related to site specific variation of the replacement patter: this suggests that the discordant topology obtained by Beast2 is likely an artefact due to model inadequacy.

**Figure 1.**
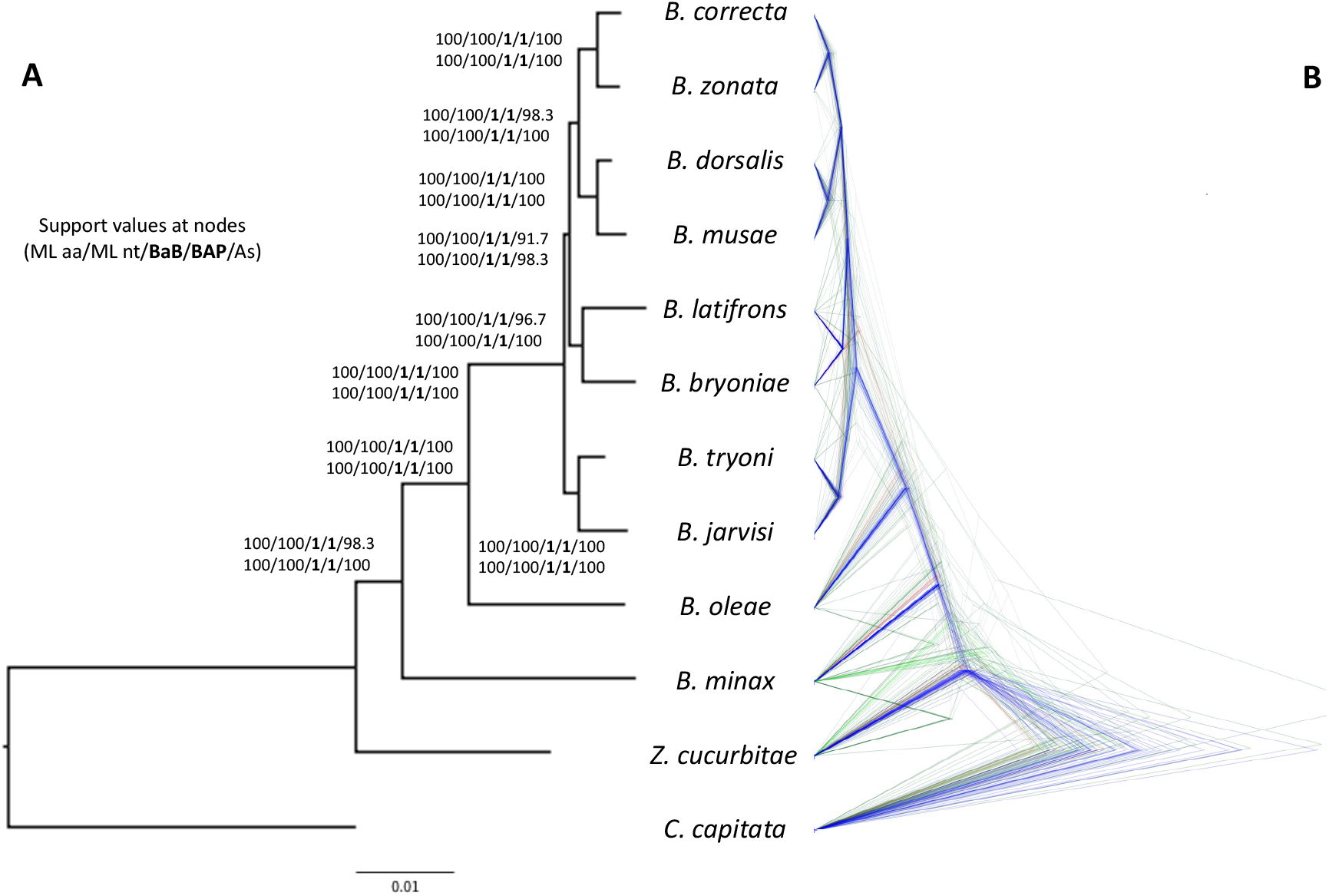
Phylogeny of *Bactrocera* inferred from the amino acidic alignments of 110 orthologous nuclear genes. A) Phylogeny obtained using a Maximum Likelihood phylogenetic analysis on the concatenated amino acidic alignments (63,297 aa). Support at nodes is given as bootstrap values for the ML analyses (both for the aminoacidic and nucleotide alignments), bootstrap values estimated by performing 100 multi-locus gene+site resampling using a multi-locus coalescent-aware phylogenetic analysis in ASTRAL across all 110 genes (As) and as posterior probabilities for the Bayesian Beast2 and PhyloBayes analyses on the amino acidic dataset (BaB and BaP, respectively). B) Bayesian analysis obtained by StarBeast2, which employs a multispecies coalescent method to estimate species trees from multiple sequence alignments (i.e., one for each of the 110 orthologous gene sets). For this analysis we used the amino acidic alignments, linking the site models across the gene sets. Note the numerous discordant gene trees, especially within the *B. dorsalis* - *B. latifrons* - *B. tryoni* clade, compared to the species tree (supported by the gene trees in blue).

Our findings are in contrast with all phylogenies inferred from mitochondrial sequence (e.g. (Yong *et al.*, 2016; A. Bin Zhang *et al.*, 2010)), which supported a closer relationship between *B. dorsalis* and *B. tryoni*. The results are instead consistent with a phylogenetic analysis of 167 Dacini species, including *Bactrocera* (San Jose *et al.*, 2018). This study, however, was not conclusive in determining the relationships between these three species, since the phylogeny had many unresolved nodes, including those relative to the most common ancestors of *B. dorsalis*, *B. latifrons* and *B. tryoni*. The low power to disentangle such relationships likely derives from the small dataset – seven nuclear genes – used to produce their phylogeny. Choo and colleagues (Choo *et al.*, 2019) analysed 116 orthologous genes across 11 *Bactrocera* species: despite the larger dataset their results are not adequately supported, as the split between (*B. dorsalis*, *B. latifrons*) and *B. tryoni* in a ML analysis had a bootstrap value of 70, indicating lack of statistical confidence.

The discordance between the results points to a complex evolutionary history of the group, as suggested by coalescent aware analyses. Combining single genes analyses into a coalescent framework also supports *B. latifrons* as the closest relative of *B. dorsalis*, both when using ASTRAL and StarBeast2 (Supplementary Figures 12-13). It is evident, however, that many genes have phylogenies not consistent with the inferred species phylogeny. For instance, the StarBeast2 approach reveals a high number of genes having an alternative phylogeny within the ((*B. dorsalis*, *B. latifrons*), *B. tryoni*), suggesting incomplete lineage sorting due to fast radiation or (ongoing) hybridization. For example, in the nucleotide datasets, 58 genes have a ((*B. dorsalis*, *B. latifrons*), *B. tryoni*) topology, 38 a ((*B. dorsalis*, *B. tryoni*), *B. latifrons*) topology and 14 a ((*B. tryoni*, *B. latifrons*), *B. dorsalis*) topology. This uncertainty is also apparent in the ASTRAL results, whereby the support for such clade falls to < 92 when bootstrapping by gene resampling (compare Supplementary Figures 8 and 9).

### Dating analysis suggests fast and recent radiation in *Bactrocera*

Our dating analysis is based on the best combination of priors according to a model selection (Supplementary Table 2) that indicated as favoured model the one where we set the mutation prior with a log-normal distribution, a strict clock, and a Birth-Death model. The Bayes Factor values, even after correcting for uncertainty by subtracting the corresponding standard deviations, are well above two, which provides overwhelming support for that model (Kass & Raftery, 1995). The fact that a strict clock is favoured over a relaxed clock is consistent with the low mean value of the coefficient of variation parameter (i.e., the standard deviation of branch rates divided by the mean rate), which equals 0.24. Therefore, we will report the results obtained by this analysis.

Incomplete lineage sorting is expected for rapid radiations (e.g., (Pollard *et al.*, 2006)., which is exactly what it is revealed by our molecular clock analyses. Consistent with a rapid radiation of the (*B. dorsalis*, *B. latifrons*, *B. tryoni*) clade, the results of the clock analysis place its origin in the mid-Pliocene, at ~2.08 mya, with a subsequent very close cladogenesis, at ~1.87 mya, separating *B. dorsalis* and *B. latifrons* (Fig. 2; see also Supplementary Figures 13-20 for the time trees obtained using the different models reported in Supplementary Table 2). Interestingly, during this period sea rose at peak levels (Zhong *et al.*, 2004) and thus increased distances between islands and island groups, possibly facilitating allopatric speciation (the three species have native ranges in south east Asia and Australia). The proximity of the two cladogenetic events and the large overlap of their 95% confidence intervals agrees with a rapid radiation, which could have resulted in frequent incomplete lineage sorting. This would also explain the discordant results between the nuclear and the mitochondrial phylogenies, a finding that is commonly reported in many organisms, including insects (Beltrán *et al.*, 2002; DeSalle & Giddings, 1986; Putnam *et al.*, 2007; Toews & Brelsford, 2012).

**Figure 2.**
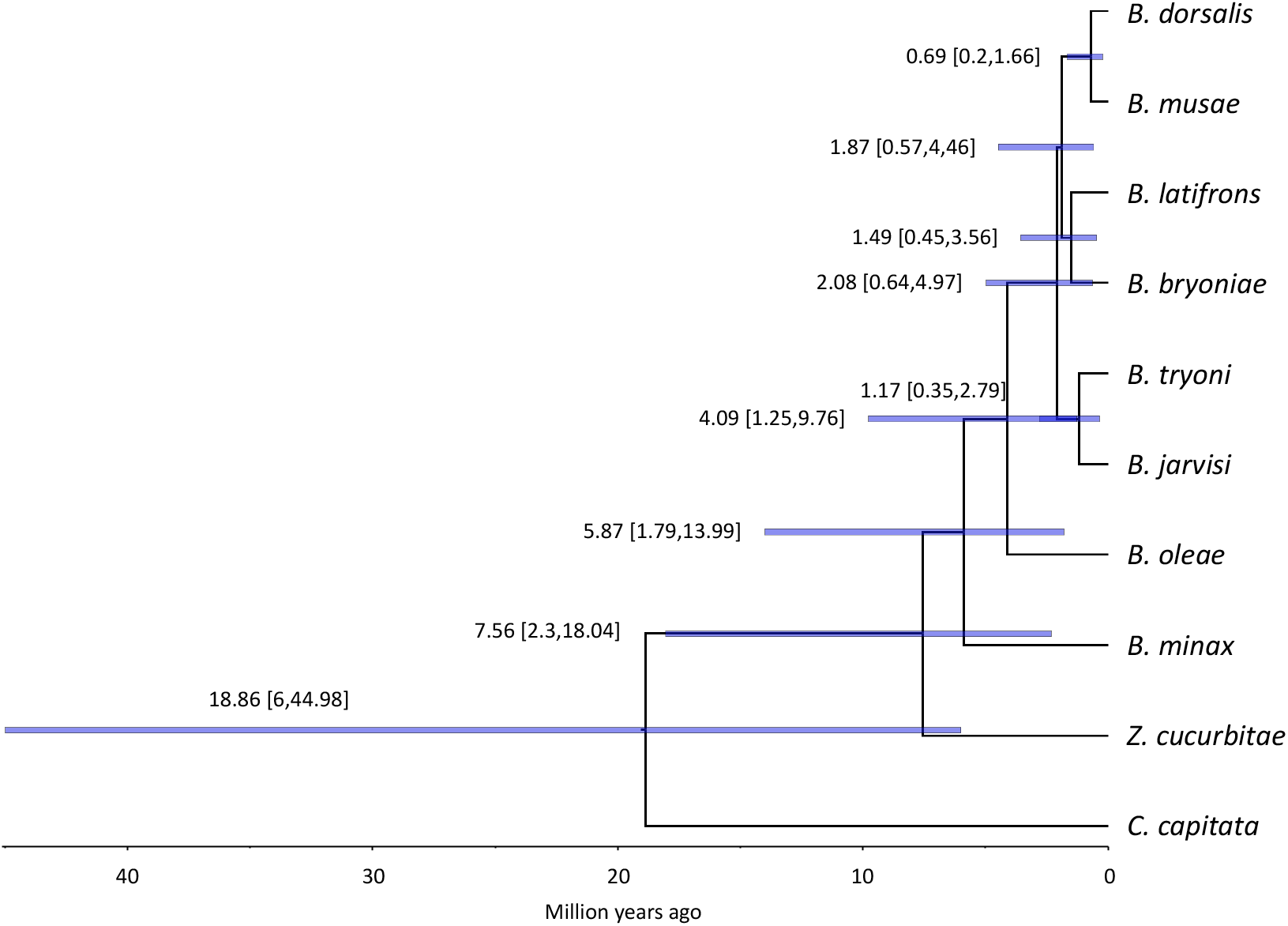
Molecular time tree of *Bactrocera* (using *C. capitata* as outgroup)*. Bactrocera* (plus *Zeugodacus*) originated during the Miocene optimum (around 19 million years ago, MYA) and experienced recent fast cladogenetic events around 2 MYA. The analysis was done setting the mutation rate log-normally distributed as prior, a strict clock and a Birth-Death model. Mean and 95% height posterior density of the inferred age (blue bars) are reported for each node.

Moreover, we cannot exclude the possibility that these species experienced, or even still experience, hybridization events, which could then result in widespread introgression events. Indeed, hybrids have been reported for several closely related *Bactrocera* species (Augustinos *et al.*, 2014; Bo *et al.*, 2014; Cruickshank *et al.*, 2001; Drew & Lambert, 1986; Ebina & Ohto, 2006; Gilchrist *et al.*, 2014; Pike *et al.*, 2003; Schutze *et al.*, 2013; Shearman *et al.*, 2010; Wee & Tan, 2005; Yeap *et al.*, 2020), and although none of the published studies involved pair of species analysed in our analyses, possible introgression can occur via direct hybridization or via intermediate hybridization events involving other closely related species.

Our results also revealed that the split between *Zeugodacus+Bactrocera* and *C. capitata* occurred less than 20 million years ago. Previously published works proposed that *C. capitata* and *Bactrocera* diverged around 24.9 mya (no confidence intervals reported, (Yaakop *et al.*, 2015)), 83 mya (95% height posterior density: 64–103 mya; (Nardi *et al.*, 2010)), 110.9 mya (95% confidence interval: 91.2–131.4 mya; (Krosch *et al.*, 2012)). The difference in the estimates could be due to several factors such as the type of molecular markers (nuclear or mitochondrial), and to the choice of priors: the clock and demographic model, the calibration points, the calibrating mutation rate, and the number of generations per year assumed. Generation time greatly influences the divergence time estimation: for example, if in our analysis we assumed five generations per year (instead of 8) without changing the other parameters, the divergence time would increase from ~20 to ~32 mya. The latter is close to what was estimated by Choo and colleagues, who performed phylogenetic and dating analyses using nuclear (*n* = 116) and mitochondrial (*n* = 13) genes of a similar species dataset (Choo *et al.*, 2019) and whose divergence time confidence intervals partially overlap with ours (31.21 (41.27–21.61 HPD) mya; this study: 18,86 (6–44.98 HPD) mya). We also employed a mutation rate prior for the four-fold degenerate sites (Obbard *et al.*, 2012) that is more conservative than the one used by Choo and colleagues. The molecular clock is known to change among sites and lineages (Drummond *et al.*, 2006), and through time (Ho *et al.*, 2011; Ho & Lo, 2013). Therefore, when choosing a rate prior it is important that it refers to a similar timescale, to sites that evolve similarly, and to species with a similar generation time. Our approach used a molecular rate estimated from *Drosophila*, which has a generation time and a phylogeny timescale similar to *Bactrocera*; hence, the four-fold degenerate site rate prior may indeed be a reliable assumption. In contrast, Choo and colleagues used a different approach, whereby they specified as prior only the divergence between *Rhagoletis* (Diptera: Tephritidae) and *Drosophila* previously inferred in (Misof *et al.*, 2014).

Finally, we would also like to point out that the mutation rate prior distribution can be a strong determinant of the divergence times estimates and therefore need to be carefully tested against alternative models (Supplementary Table 2): for example, mean divergence time between *C. capitata* and *Bactrocera* is estimated between 18.3 and 18.9 mya when using a log-normal distribution (Fig. 2 and Supplementary Figures 14-16), while it is estimated between 13 and 13.2 mya using a normal distribution (Supplementary Figures 17-21), and the same reduction of ~30% holds for all other divergence times across the phylogeny. The confidence intervals are overlapping and therefore such means are not significantly different, however model selection indicates that a log-normal as the most fitting prior distribution of the rate.

Overall, our results highlight the importance of using genome-wide data to resolve complex phylogenies and provide a useful framework for future comparative genomics and comparative biology studies in *Bactrocera*. The possibility that hybridization can still occur between closely related species also warns about the possibility that selective events in one species (for instance, resistance to insecticides) may be readily transferred to other species by introgression.

## References

Abascal, F., Zardoya, R., & Telford, M. J. (2010). TranslatorX: Multiple alignment of nucleotide sequences guided by amino acid translations. Nucleic Acids Research, 38(SUPPL. 2), W7–13. https://doi.org/10.1093/nar/gkq291

Augustinos, A. A., Drosopoulou, E., Gariou-Papalexiou, A., Bourtzis, K., Mavragani-Tsipidou, P., & Zacharopoulou, A. (2014). The Bactrocera dorsalis species complex: Comparative cytogenetic analysis in support of Sterile Insect Technique applications. BMC Genetics, 15(2), S16. https://doi.org/10.1186/1471-2156-15-S2-S16

Barido-Sottani, J., Bošková, V., Plessis, L. Du, Kühnert, D., Magnus, C., Mitov, V., Müller, N. F., Pečerska, J., Rasmussen, D. A., Zhang, C., Drummond, A. J., Heath, T. A., Pybus, O. G., Vaughan, T. G., & Stadler, T. (2018). Taming the BEAST - A Community Teaching Material Resource for BEAST 2. Systematic Biology, 67(1), 170–174. https://doi.org/10.1093/sysbio/syx060

Bayega, A., Djambazian, H., Tsoumani, K. T., Gregoriou, M. E., Sagri, E., Drosopoulou, E., Mavragani-Tsipidou, P., Giorda, K., Tsiamis, G., Bourtzis, K., Oikonomopoulos, S., Dewar, K., Church, D. M., Papanicolaou, A., Mathiopoulos, K. D., & Ragoussis, J. (2020). De novo assembly of the olive fruit fly (Bactrocera oleae) genome with linked-reads and long-read technologies minimizes gaps and provides exceptional y chromosome assembly. BMC Genomics, 21(1). https://doi.org/10.1186/s12864-020-6672-3

Beltrán, M., Jiggins, C. D., Bull, V., Linares, M., Mallet, J., McMillan, W. O., & Bermingham, E. (2002). Phylogenetic Discordance at the Species Boundary: Comparative Gene Genealogies Among Rapidly Radiating Heliconius Butterflies. Molecular Biology and Evolution, 19(12), 2176–2190. https://doi.org/10.1093/oxfordjournals.molbev.a004042

Bickel, D., Pape, T., & Meier, R. (2009). Diptera Diversity: Status, Challenges and Tools. Brill. https://doi.org/10.1163/ej.9789004148970.I-459

Bo, W., Ahmad, S., Dammalage, T., Tomas, U. S., Wornoayporn, V., Ul Haq, I., Cáceres, C., Vreysen, M. J. B., & Schutze, M. K. (2014). Mating compatibility between Bactrocera invadens and Bactrocera dorsalis (Diptera: Tephritidae). Journal of Economic Entomology, 107(2), 623–629. https://doi.org/10.1603/ec13514

Bouckaert, R., Vaughan, T. G., Barido-Sottani, J., Duchêne, S., Fourment, M., Gavryushkina, A., Heled, J., Jones, G., Kühnert, D., De Maio, N., Matschiner, M., Mendes, F. K., Müller, N. F., Ogilvie, H. A., Du Plessis, L., Popinga, A., Rambaut, A., Rasmussen, D., Siveroni, I., … Drummond, A. J. (2019). BEAST 2.5: An advanced software platform for Bayesian evolutionary analysis. PLoS Computational Biology, 15(4), e1006650. https://doi.org/10.1371/journal.pcbi.1006650

Camacho, C., Coulouris, G., Avagyan, V., Ma, N., Papadopoulos, J., Bealer, K., & Madden, T. L. (2009). BLAST+: Architecture and applications. BMC Bioinformatics, 10(1), 421. https://doi.org/10.1186/1471-2105-10-421

Choo, A., Nguyen, T. N. M., Ward, C. M., Chen, I. Y., Sved, J., Shearman, D., Gilchrist, A. S., Crisp, P., & Baxter, S. W. (2019). Identification of Y-chromosome scaffolds of the Queensland fruit fly reveals a duplicated gyf gene paralogue common to many Bactrocera pest species. Insect Molecular Biology, 28(6), 873–886. https://doi.org/10.1111/imb.12602

Cruickshank, L., Jessup, A. J., & Cruickshank, D. J. (2001). Interspecific crosses of Bactrocera tryoni (Froggatt) and Bactrocera jarvisi (Tryon) (Diptera: Tephritidae) in the laboratory. Australian Journal of Entomology, 40(3), 278–280. https://doi.org/10.1046/j.1440-6055.2001.00223.x

Daane, K. M., & Johnson, M. W. (2010). Olive fruit fly: Managing an ancient pest in modern times. Annual Review of Entomology, 55, 151–169. https://doi.org/10.1146/annurev.ento.54.110807.090553

DeSalle, R., & Giddings, L. V. (1986). Discordance of nuclear and mitochondrial DNA phylogenies in Hawaiian Drosophila. Proceedings of the National Academy of Sciences, 83(18), 6902–6906. https://doi.org/10.1073/pnas.83.18.6902

Drew, R. A. I. (1972). The Generic and Subgeneric Classification of Dacini (diptera: Tephritidae) from the South Pacific Area. Australian Journal of Entomology, 11(1), 1–22. https://doi.org/10.1111/j.1440-6055.1972.tb01601.x

Drew, R. A. I. (1989). The tropical fruit flies (Diptera: Tephritidae: Dacinae) of the Australasian region and Oceanic regions. Memoirs of the Queensland Museum, 26, 1–521.

Drew, R. A. I., & Hancock, D. L. (1999). Phylogeny of the Tribe Dacini (Dacinae) Based on Morphological, Distributional, and Biological Data. In Fruit Flies (Tephritidae) (pp. 509–522). CRC Press. https://doi.org/10.1201/9781420074468-28

Drew, R. A. I., & Lambert, D. M. (1986). On the Specific Status of Dacus (Bactrocera) aquilonis and D. (Bactrocera) tryoni (Diptera: Tephritidae). Annals of the Entomological Society of America, 79(6), 870–878. https://doi.org/10.1093/aesa/79.6.870

Drew, R. A. I., & Romig, M. C. (2016). Keys to the Tropical Fruit Flies of South-East Asia. CABI.

Drummond, A. J., Ho, S. Y. W., Phillips, M. J., & Rambaut, A. (2006). Relaxed Phylogenetics and Dating with Confidence. PLOS Biology, 4(5), e88. https://doi.org/10.1371/journal.pbio.0040088

Ebina, T., & Ohto, K. (2006). Morphological characters and PCR-RFLP markers in the interspecific hybrids between Bactrocera carambolae and B. papayae of the B. dorsalis species complex (Diptera: Tephritidae). Res. Bull. Plant Prot. Japan, 42, 23–34.

Geib, S. M., Calla, B., Hall, B., Hou, S., & Manoukis, N. C. (2014). Characterizing the developmental transcriptome of the oriental fruit fly, Bactrocera dorsalis (Diptera: Tephritidae) through comparative genomic analysis with Drosophila melanogaster utilizing modENCODE datasets. BMC Genomics, 15(1). https://doi.org/10.1186/1471-2164-15-942

Gilchrist, A. S., Shearman, D. C., Frommer, M., Raphael, K. A., Deshpande, N. P., Wilkins, M. R., Sherwin, W. B., & Sved, J. A. (2014). The draft genome of the pest tephritid fruit fly Bactrocera tryoni: Resources for the genomic analysis of hybridising species. BMC Genomics, 15(1), 1153. https://doi.org/10.1186/1471-2164-15-1153

Grabherr, M. G., Haas, B. J., Yassour, M., Levin, J. Z., Thompson, D. A., Amit, I., Adiconis, X., Fan, L., Raychowdhury, R., Zeng, Q., Chen, Z., Mauceli, E., Hacohen, N., Gnirke, A., Rhind, N., di Palma, F., Birren, B. W., Nusbaum, C., Lindblad-Toh, K., … Regev, A. (2011). Full-length transcriptome assembly from RNA-Seq data without a reference genome. Nature Biotechnology, 29(7), 644–652. https://doi.org/10.1038/nbt.1883

Ho, S. Y. W., Lanfear, R., Bromham, L., Phillips, M. J., Soubrier, J., Rodrigo, A. G., & Cooper, A. (2011). Time-dependent rates of molecular evolution. Molecular Ecology, 20(15), 3087–3101. https://doi.org/10.1111/j.1365-294X.2011.05178.x

Ho, S. Y. W., & Lo, N. (2013). The insect molecular clock. Australian Journal of Entomology, 52(2), 101–105. https://doi.org/10.1111/aen.12018

Junqueira, A. C. M., Azeredo-Espin, A. M. L., Paulo, D. F., Marinho, M. A. T., Tomsho, L. P., Drautz-Moses, D. I., Purbojati, R. W., Ratan, A., & Schuster, S. C. (2016). Large-scale mitogenomics enables insights into Schizophora (Diptera) radiation and population diversity. Scientific Reports, 6(1), 1–13. https://doi.org/10.1038/srep21762

Kass, R. E., & Raftery, A. E. (1995). Bayes factors. Journal of the American Statistical Association. https://doi.org/10.1080/01621459.1995.10476572

Katoh, K., & Standley, D. M. (2013). MAFFT multiple sequence alignment software version 7: Improvements in performance and usability. Molecular Biology and Evolution, 30(4), 772–780. https://doi.org/10.1093/molbev/mst010

Krosch, M. N., Schutze, M. K., Armstrong, K. F., Graham, G. C., Yeates, D. K., & Clarke, A. R. (2012). A molecular phylogeny for the Tribe Dacini (Diptera: Tephritidae): Systematic and biogeographic implications. Molecular Phylogenetics and Evolution, 64(3), 513–523. https://doi.org/10.1016/j.ympev.2012.05.006

Lartillot, N., & Philippe, H. (2004). A Bayesian Mixture Model for Across-Site Heterogeneities in the Amino-Acid Replacement Process. Molecular Biology and Evolution, 21(6), 1095–1109. https://doi.org/10.1093/molbev/msh112

Li, X., Yang, H., Wang, T., Wang, J., & Wei, H. (2019). Life history and adult dynamics of Bactrocera dorsalis in the citrus orchard of Nanchang, a subtropical area from China: Implications for a control timeline. ScienceAsia, 45(3), 212–220. https://doi.org/10.2306/scienceasia1513-1874.2019.45.212

Löytynoja, A., & Goldman, N. (2008). A model of evolution and structure for multiple sequence alignment. Philosophical Transactions of the Royal Society B: Biological Sciences, 363(1512), 3913–3919. https://doi.org/10.1098/rstb.2008.0170

Misof, B., Liu, S., Meusemann, K., Peters, R. S., Donath, A., Mayer, C., Frandsen, P. B., Ware, J., Flouri, T., Beutel, R. G., Niehuis, O., Petersen, M., Izquierdo-Carrasco, F., Wappler, T., Rust, J., Aberer, A. J., Aspock, U., Aspock, H., Bartel, D., … Zhou, X. (2014). Phylogenomics resolves the timing and pattern of insect evolution. Science, 346(6210), 763–767. https://doi.org/10.1126/science.1257570

Muraji, M., & Nakahara, S. (2001). Phylogenetic relationships among fruit flies, Bactrocera (Diptera, Tephritidae), based on the mitochondrial rDNA sequences. Insect Molecular Biology, 10(6), 549–559. https://doi.org/10.1046/j.0962-1075.2001.00294.x

Nakahara, S., & Muraji, M. (2007). Phylogenetic Analyses of Bactrocera Fruit Flies (Diptera: Tephritidae) Based on Nucleotide Sequences of the Mitochondrial COI and COII Genes. Research Bulletin Plant Protection Japan, 44.

Nardi, F., Carapelli, A., Boore, J. L., Roderick, G. K., Dallai, R., & Frati, F. (2010). Domestication of olive fly through a multi-regional host shift to cultivated olives: Comparative dating using complete mitochondrial genomes. Molecular Phylogenetics and Evolution, 57(2), 678–686. https://doi.org/10.1016/j.ympev.2010.08.008

Norrbom, A. (1994). New genera of Tephritidae (Diptera) from Brazil and Dominican Amber, with phylogenetic analysis of the tribe Ortalotrypetini. Insecta Mundi, 8(1–2), 1–15.

Obbard, D. J., MacLennan, J., Kim, K. W., Rambaut, A., O’Grady, P. M., & Jiggins, F. M. (2012). Estimating divergence dates and substitution rates in the drosophila phylogeny. Molecular Biology and Evolution, 29(11), 3459–3473. https://doi.org/10.1093/molbev/mss150

Ogilvie, H. A., Bouckaert, R. R., & Drummond, A. J. (2017). StarBEAST2 brings faster species tree inference and accurate estimates of substitution rates. Molecular Biology and Evolution, 34(8), 2101–2114. https://doi.org/10.1093/molbev/msx126

Ometto, L., Cestaro, A., Ramasamy, S., Grassi, A., Revadi, S., Siozios, S., Moretto, M., Fontana, P., Varotto, C., Pisani, D., Dekker, T., Wrobel, N., Viola, R., Pertot, I., Cavalieri, D., Blaxter, M., Anfora, G., & Rota-Stabelli, O. (2013). Linking Genomics and Ecology to Investigate the Complex Evolution of an Invasive Drosophila Pest. Genome Biology and Evolution, 5(4), 745–757. https://doi.org/10.1093/gbe/evt034

Papanicolaou, A., Schetelig, M. F., Arensburger, P., Atkinson, P. W., Benoit, J. B., Bourtzis, K., Castañera, P., Cavanaugh, J. P., Chao, H., Childers, C., Curril, I., Dinh, H., Doddapaneni, H., Dolan, A., Dugan, S., Friedrich, M., Gasperi, G., Geib, S., Georgakilas, G., … Handler, A. M. (2016). The whole genome sequence of the Mediterranean fruit fly, Ceratitis capitata (Wiedemann), reveals insights into the biology and adaptive evolution of a highly invasive pest species. Genome Biology, 17(1), 192. https://doi.org/10.1186/s13059-016-1049-2

Pike, N., Wang, W. Y. S., & Meats, A. (2003). The likely fate of hybrids of Bactrocera tryoni and Bactrocera neohumeralis. Heredity, 90(5), 365–370. https://doi.org/10.1038/sj.hdy.6800253

Pollard, D. A., Iyer, V. N., Moses, A. M., & Eisen, M. B. (2006). Widespread Discordance of Gene Trees with Species Tree in Drosophila: Evidence for Incomplete Lineage Sorting. PLoS Genetics, 2(10), e173. https://doi.org/10.1371/journal.pgen.0020173

Putnam, A. S., Scriber, J. M., & Andolfatto, P. (2007). Discordant divergence times among Z-chromosome regions between two ecologically distinct swallowtail butterfly species. Evolution, 61(4), 912–927. https://doi.org/10.1111/j.1558-5646.2007.00076.x

Ramasamy, S., Ometto, L., Crava, C. M., Revadi, S., Kaur, R., Horner, D. S., Pisani, D., Dekker, T., Anfora, G., & Rota-Stabelli, O. (2016). The evolution of olfactory gene families in Drosophila and the genomic basis of chemical-ecological adaptation in Drosophila suzukii. Genome Biology and Evolution, 8(8), 2297–2311. https://doi.org/10.1093/gbe/evw160

Russel, P. M., Brewer, B. J., Klaere, S., & Bouckaert, R. R. (2019). Model Selection and Parameter Inference in Phylogenetics Using Nested Sampling. Systematic Biology, 68(2), 219–233. https://doi.org/10.1093/sysbio/syy050

San Jose, M., Doorenweerd, C., Leblanc, L., Barr, N., Geib, S., & Rubinoff, D. (2018). Incongruence between molecules and morphology: A seven-gene phylogeny of Dacini fruit flies paves the way for reclassification (Diptera: Tephritidae). Molecular Phylogenetics and Evolution, 121, 139–149. https://doi.org/10.1016/j.ympev.2017.12.001

Schutze, M. K., Jessup, A., Ul-Haq, I., Vreysen, M. J. B., Wornoayporn, V., Vera, M. T., & Clarke, A. R. (2013). Mating Compatibility Among Four Pest Members of the Bactrocera dorsalis Fruit Fly Species Complex (Diptera: Tephritidae). Journal of Economic Entomology, 106(2), 695–707. https://doi.org/10.1603/EC12409

Shearman, D. C. A., Frommer, M., Morrow, J. L., Raphael, K. A., & Gilchrist, A. S. (2010). Interspecific Hybridization as a Source of Novel Genetic Markers for the Sterile Insect Technique in Bactrocera tryoni (Diptera: Tephritidae). Journal of Economic Entomology, 103(4), 1071–1079. https://doi.org/10.1603/EC09241

Sim, S. B., & Geib, S. M. (2017). A Chromosome-Scale Assembly of the Bactrocera cucurbitae Genome Provides Insight to the Genetic Basis of white pupae. G3: Genes, Genomes, Genetics, 7(6), 1927–1940. https://doi.org/10.1534/g3.117.040170

Smith, P. T., Kambhampati, S., & Armstrong, K. A. (2003). Phylogenetic relationships among Bactrocera species (Diptera: Tephritidae) inferred from mitochondrial DNA sequences. Molecular Phylogenetics and Evolution, 26(1), 8–17. https://doi.org/10.1016/s1055-7903(02)00293-2

Stamatakis, A. (2014). RAxML version 8: A tool for phylogenetic analysis and post-analysis of large phylogenies. Bioinformatics, 30(9), 1312–1313. https://doi.org/10.1093/bioinformatics/btu033

Stephens, A. E. A., Kriticos, D. J., & Leriche, A. (2007). The current and future potential geographical distribution of the oriental fruit fly, Bactrocera dorsalis (Diptera: Tephritidae). Bulletin of Entomological Research, 97(4), 369–378. https://doi.org/10.1017/S0007485307005044

Theron, C. D., Manrakhan, A., & Weldon, C. W. (2017). Host use of the oriental fruit fly, *Bactrocera dorsalis* (Hendel) (Diptera: Tephritidae), in South Africa. Journal of Applied Entomology, 141(10), 810–816. https://doi.org/10.1111/jen.12400

Toews, D. P. L., & Brelsford, A. (2012). The biogeography of mitochondrial and nuclear discordance in animals. Molecular Ecology, 21(16), 3907–3930. https://doi.org/10.1111/j.1365-294X.2012.05664.x

Vargas, R. I., Walsh, W. A., Kanehisa, D., Jang, E. B., & Armstrong, J. W. (1997). Demography of four Hawaiian fruit flies (diptera: Tephritidae) reared at five constant temperatures. Annals of the Entomological Society of America, 90(2), 162–168. https://doi.org/10.1093/aesa/90.2.162

Virgilio, M., Jordaens, K., Verwimp, C., White, I. M., & De Meyer, M. (2015). Higher phylogeny of frugivorous flies (Diptera, Tephritidae, Dacini): Localised partition conflicts and a novel generic classification. Molecular Phylogenetics and Evolution, 85, 171–179. https://doi.org/10.1016/j.ympev.2015.01.007

Wang, J., Xiong, K.-C., & Liu, Y.-H. (2016). De novo Transcriptome Analysis of Chinese Citrus Fly, Bactrocera minax (Diptera: Tephritidae), by High-Throughput Illumina Sequencing. PLOS ONE, 11(6), e0157656. https://doi.org/10.1371/journal.pone.0157656

Wee, S.-L., & Tan, K.-H. (2005). Evidence of natural hybridization between two sympatric sibling species of Bactrocera dorsalis complex based on pheromone analysis. Journal of Chemical Ecology, 31(4), 845–858. https://doi.org/10.1007/s10886-005-3548-6

White, I. M., & Elson-Harris, M. M. (1992). Fruit Flies of Economic Significance: Their Identification and Bionomics. CAB International, Wallingford, UK.

Wiegmann, B. M., Trautwein, M. D., Winkler, I. S., Barr, N. B., Kim, J.-W., Lambkin, C., Bertone, M. A., Cassel, B. K., Bayless, K. M., Heimberg, A. M., Wheeler, B. M., Peterson, K. J., Pape, T., Sinclair, B. J., Skevington, J. H., Blagoderov, V., Caravas, J., Kutty, S. N., Schmidt-Ott, U., … Yeates, D. K. (2011). Episodic radiations in the fly tree of life. Proceedings of the National Academy of Sciences of the United States of America, 108(14), 5690–5695. https://doi.org/10.1073/pnas.1012675108

Yaakop, S., Ibrahim, N. J., Shariff, S., & Zain, B. M. M. (2015). Molecular clock analysis on five Bactrocera species flies (Diptera: Tephritidae) based on combination of COI and NADH sequences. Oriental Insects, 49(1–2), 150–164. https://doi.org/10.1080/00305316.2015.1081421

Yeap, H. L., Lee, S. F., Robinson, F., Mourant, R. G., Sved, J. A., Frommer, M., Papanicolaou, A., Edwards, O. R., & Oakeshott, J. G. (2020). Separating two tightly linked species-defining phenotypes in Bactrocera with hybrid recombinant analysis. BMC Genetics, 21(2), 132. https://doi.org/10.1186/s12863-020-00936-1

Yong, H.-S., Song, S.-L., Lim, P.-E., Eamsobhana, P., & Suana, I. W. (2016). Complete Mitochondrial Genome of Three Bactrocera Fruit Flies of Subgenus Bactrocera (Diptera: Tephritidae) and Their Phylogenetic Implications. PLOS ONE, 11(2), e0148201. https://doi.org/10.1371/journal.pone.0148201

Zhang, A. Bin, Liu, Y. H., Wu, W. X., & Wang, Z. Le. (2010). Molecular Phylogeny of Bactrocera Species (Diptera: Tephritidae: Dacini) Inferred from Mitochondrial Sequences of 16S rDNA and COI Sequences. Florida Entomologist, 93(3), 369. https://doi.org/10.1653/024.093.0308

Zhang, C., Rabiee, M., Sayyari, E., & Mirarab, S. (2018). ASTRAL-III: Polynomial time species tree reconstruction from partially resolved gene trees. BMC Bioinformatics, 19(S6), 153. https://doi.org/10.1186/s12859-018-2129-y

Zhong, G., Geng, J., Wong, H. K., Ma, Z., & Wu, N. (2004). A semi-quantitative method for the reconstruction of eustatic sea level history from seismic profiles and its application to the southern South China Sea. Earth and Planetary Science Letters, 223(3), 443–459. https://doi.org/10.1016/j.epsl.2004.04.039

