## Supplemental Figures for "The impact of fast radiation on the phylogeny of *Bactrocera* fruit flies"

**Supplemental Figure 1.** Phylogeny of *Bactrocera* obtained by RAxML using a Maximum Likelihood analysis on the concatenated codon alignments of 110 orthologous nuclear genes (189,891 nt). Support at nodes is given as bootstrap values.

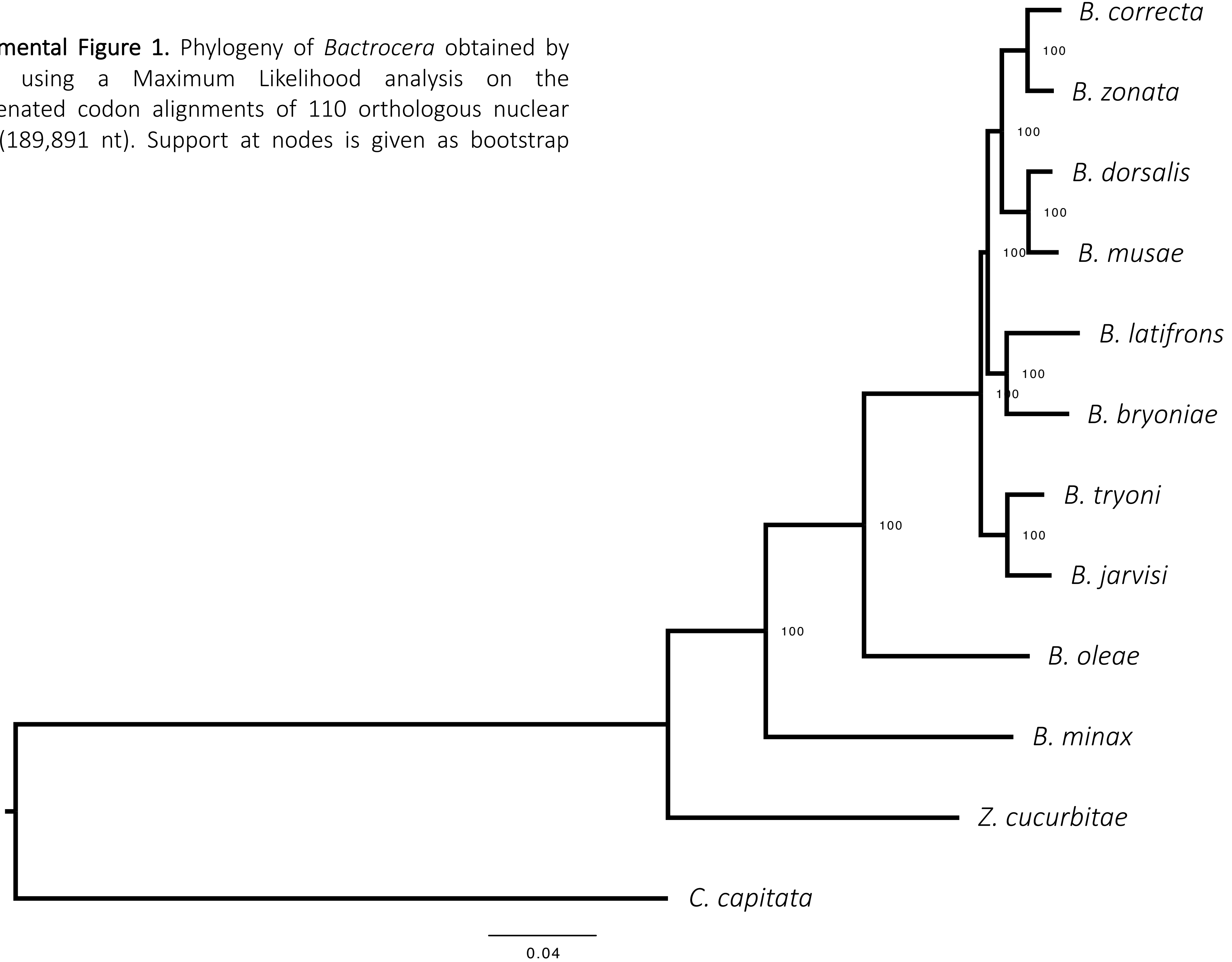

**Supplemental Figure 2.** Phylogeny of *Bactrocera* obtained by Beast2 using a Bayesian analysis on the concatenated amino acidic alignments (63,297 aa). Support at nodes is given as posterior probabilities.

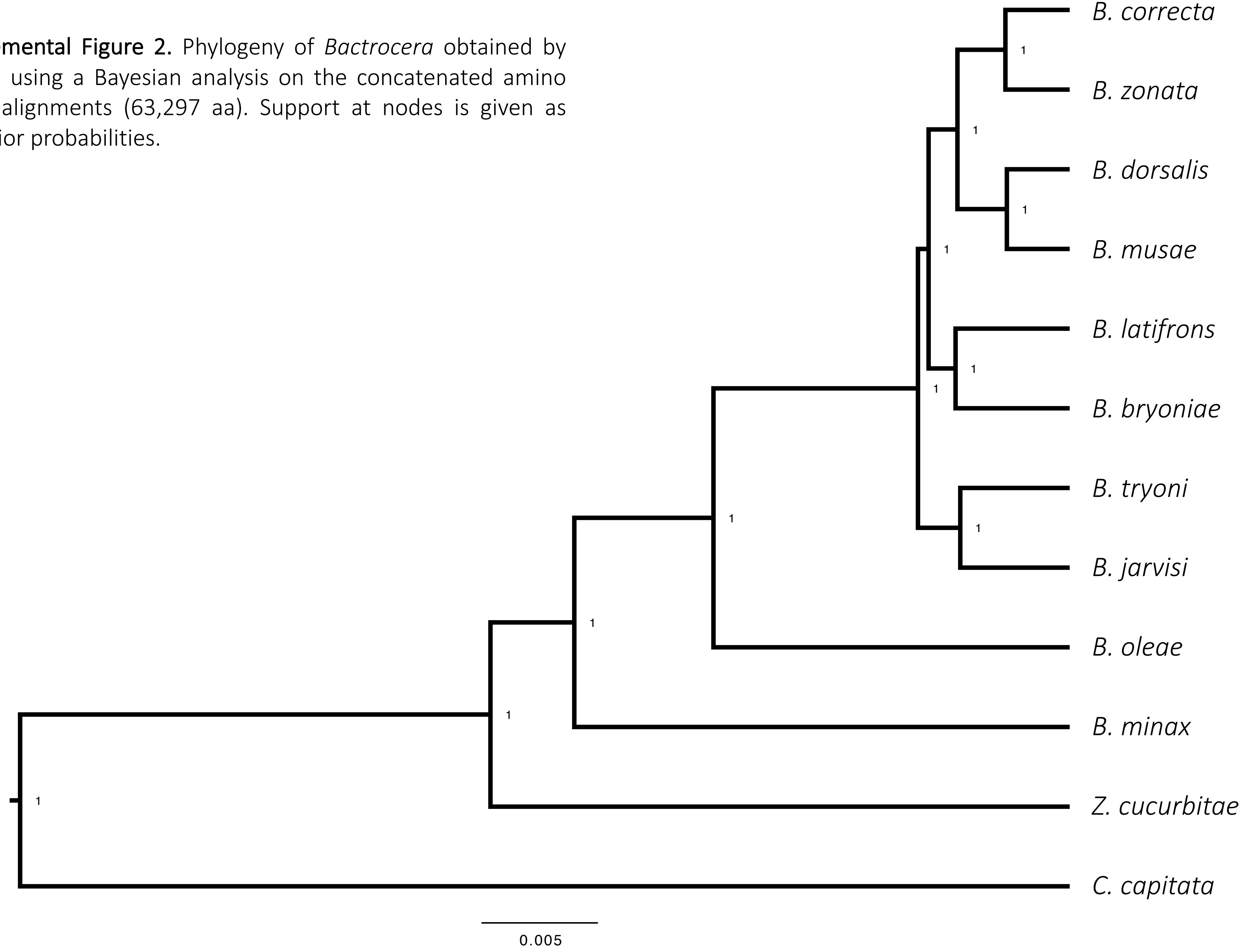

**Supplemental Figure 3.** Phylogeny of *Bactrocera* obtained by Beast2 using a Bayesian analysis on the concatenated codon alignments of 110 orthologous nuclear genes (189,891 nt). Support at nodes is given as posterior probabilities.

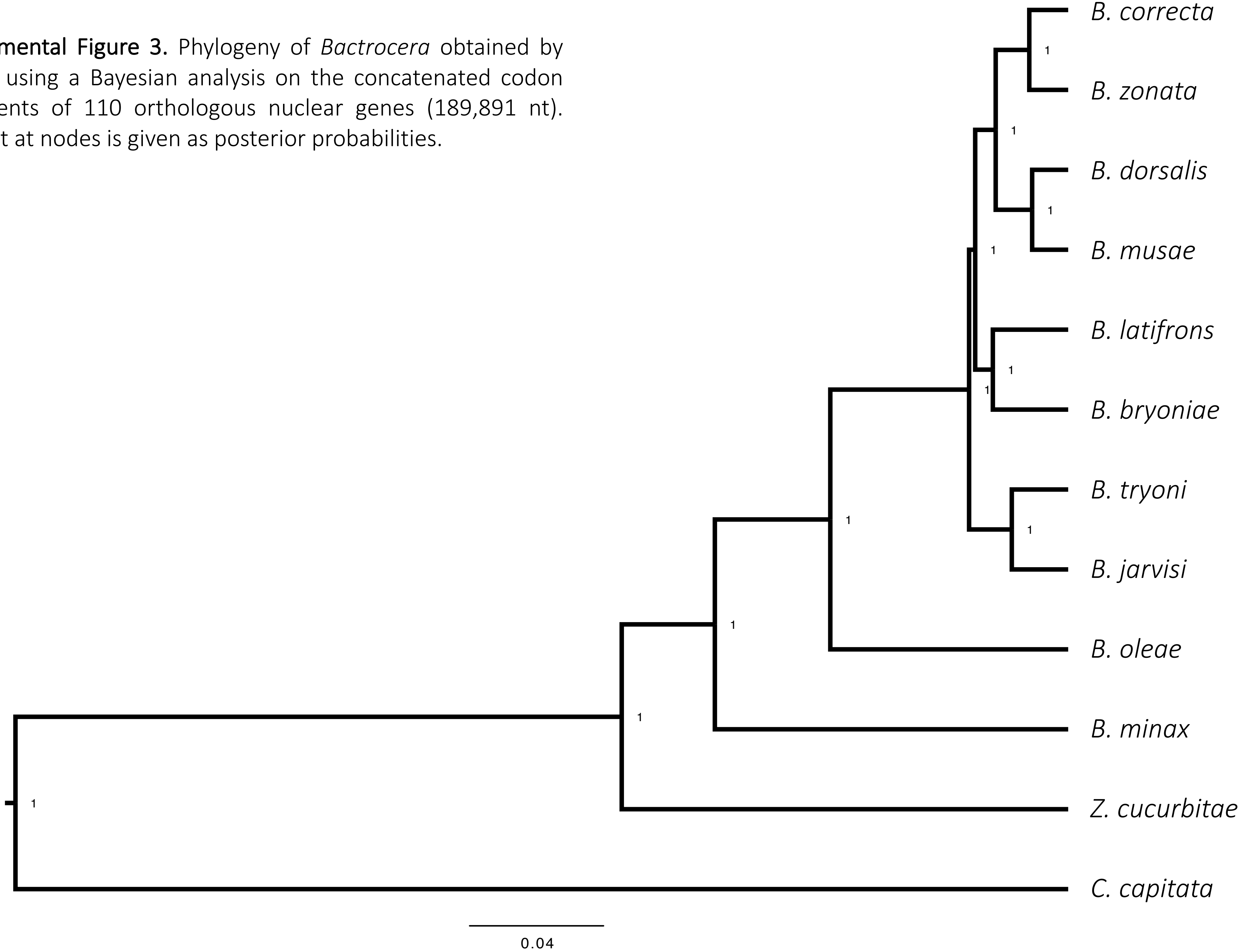

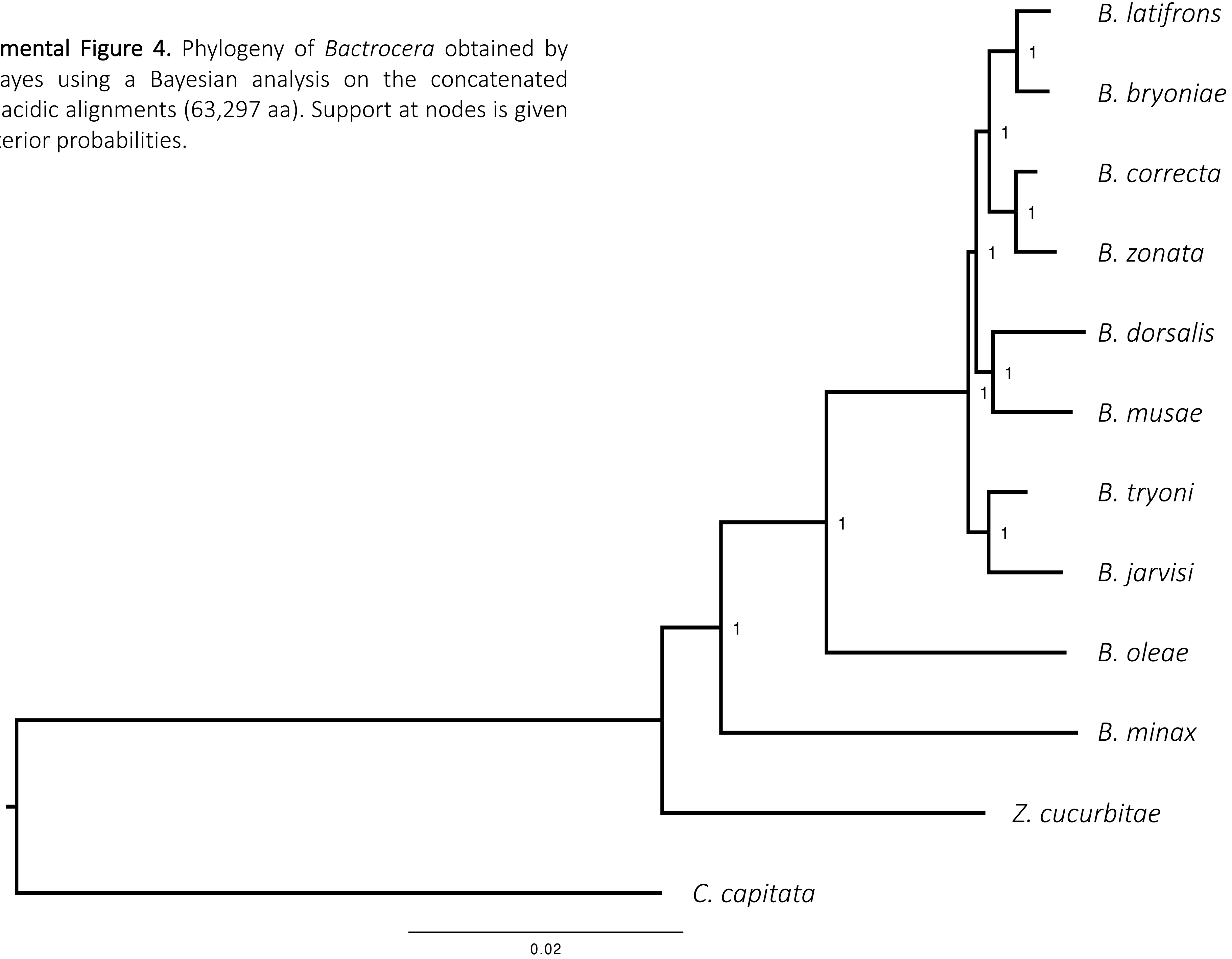


**Supplemental Figure 4.** Phylogeny of *Bactrocera* obtained by PhyloBayes using a Bayesian analysis on the concatenated amino acidic alignments (63,297 aa). Support at nodes is given as posterior probabilities.

**Supplemental Figure 5.** Phylogeny of *Bactrocera* obtained by Beast2 using a Bayesian analysis on the 4-fold degenerate sites of the concatenated alignments of 110 orthologous nuclear genes (24,885 nt). Support at nodes is given as posterior probabilities.

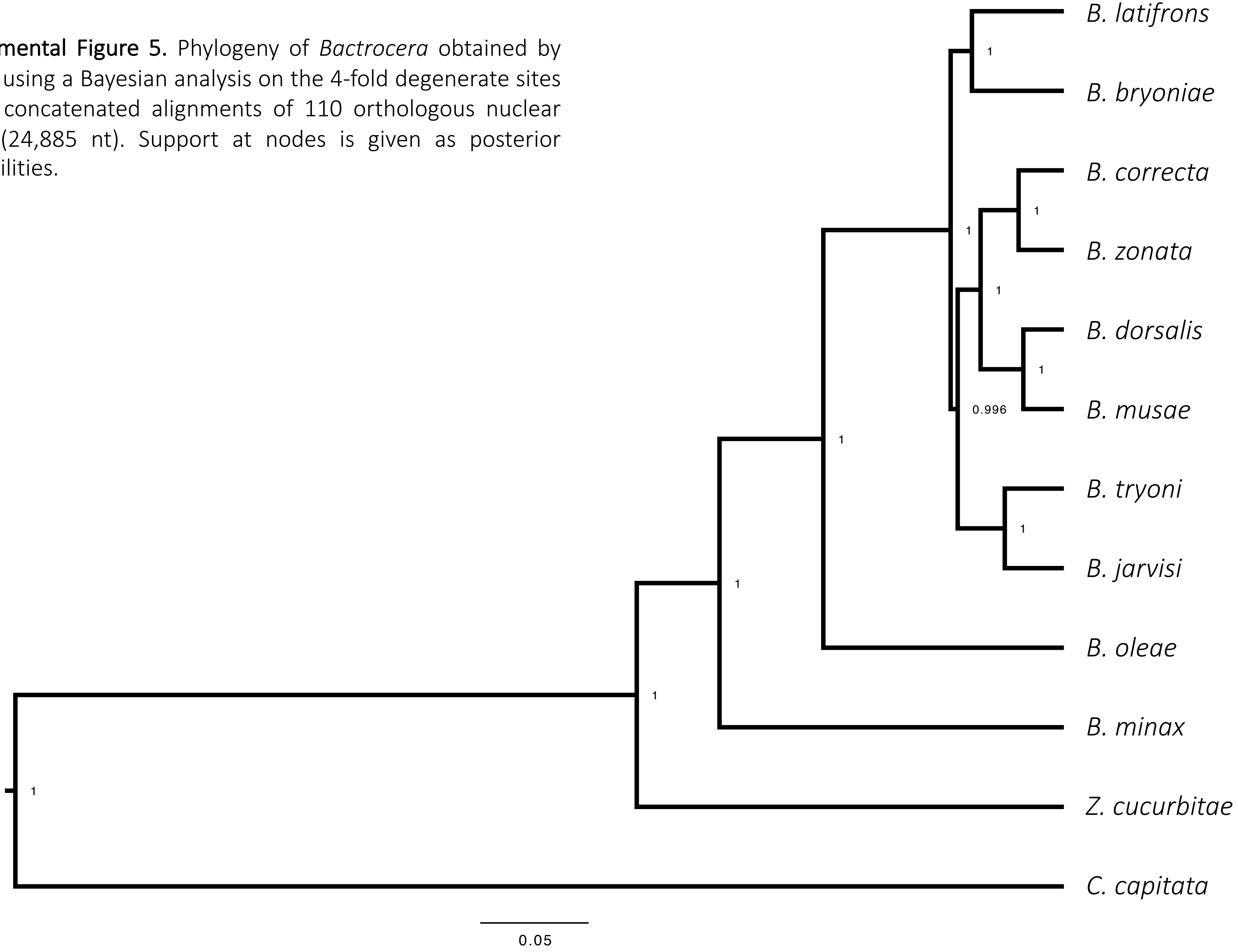

**Supplemental Figure 6.** Phylogeny of *Bactrocera* obtained by PhyloBayes using a Bayesian analysis on the 4-fold degenerate sites of the concatenated alignments of 110 orthologous nuclear genes (24,885 nt). Support at nodes is given as posterior probabilities.

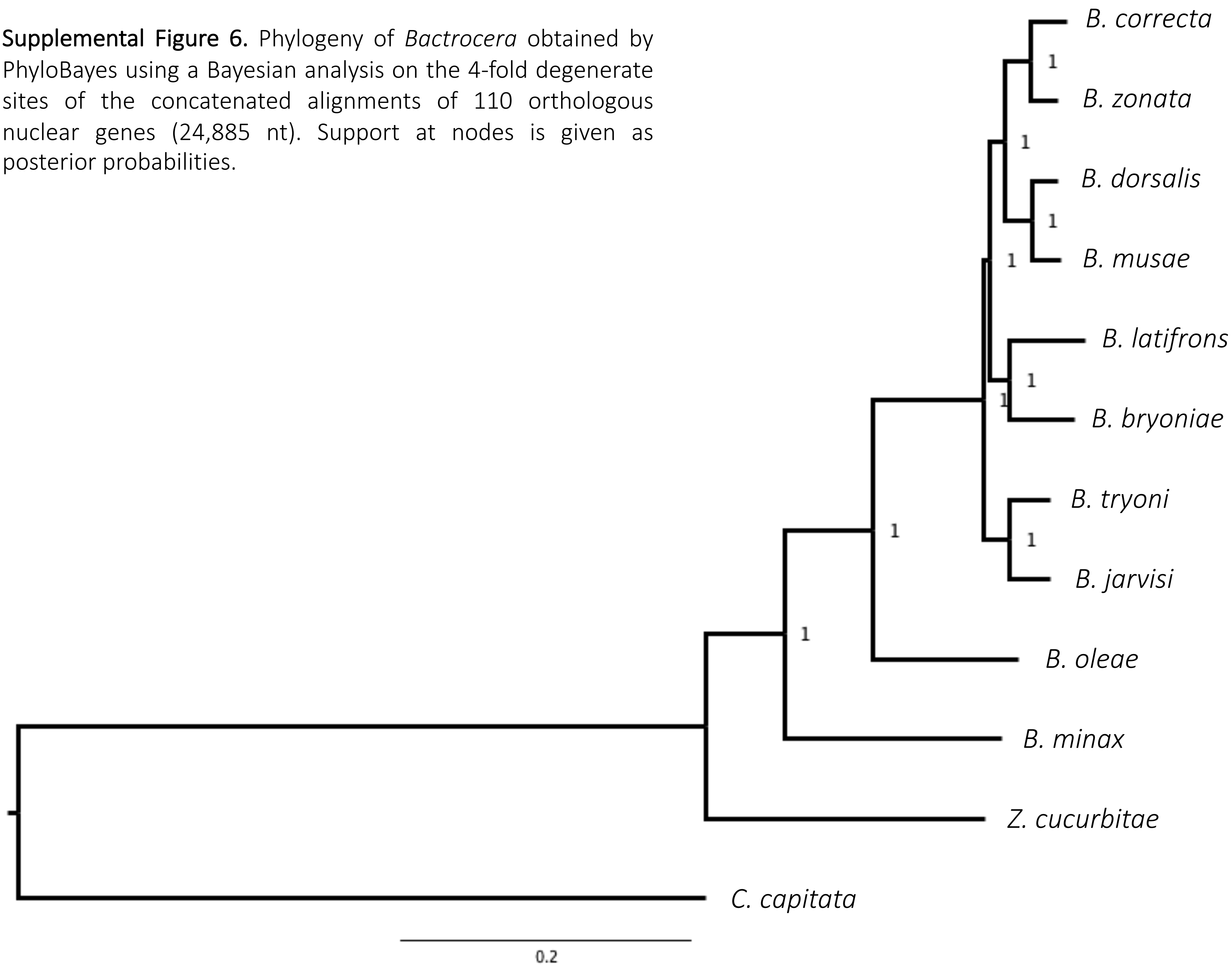

**Supplemental Figure 7.** Phylogeny of *Bactrocera* obtained by RAxML using a Maximum Likelihood analysis on the 4-fold degenerate sites of the concatenated alignments of 110 orthologous nuclear genes (24,885 nt). Support at nodes is given as bootstrap values.

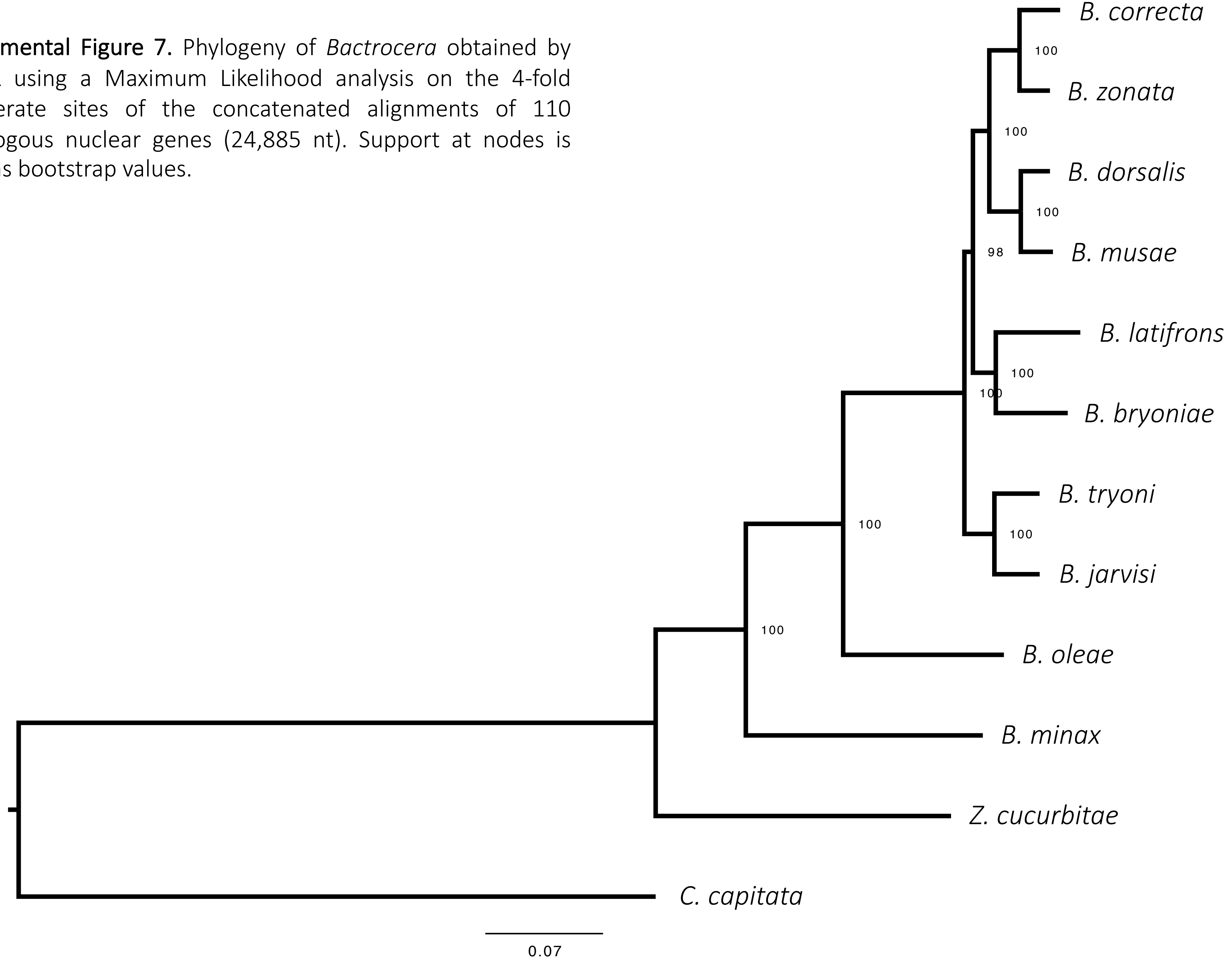

**Supplemental Figure 8.** Multi-locus coalescent-aware phylogenesis of *Bactrocera* inferred from 110 orthologous nuclear genes using ASTRAL. Analyses are based on all single ML gene trees obtained by RAxML using the aminoacidic alignments. Bootstrap values were estimated by performing 100 multi-locus bootstrap replicates.

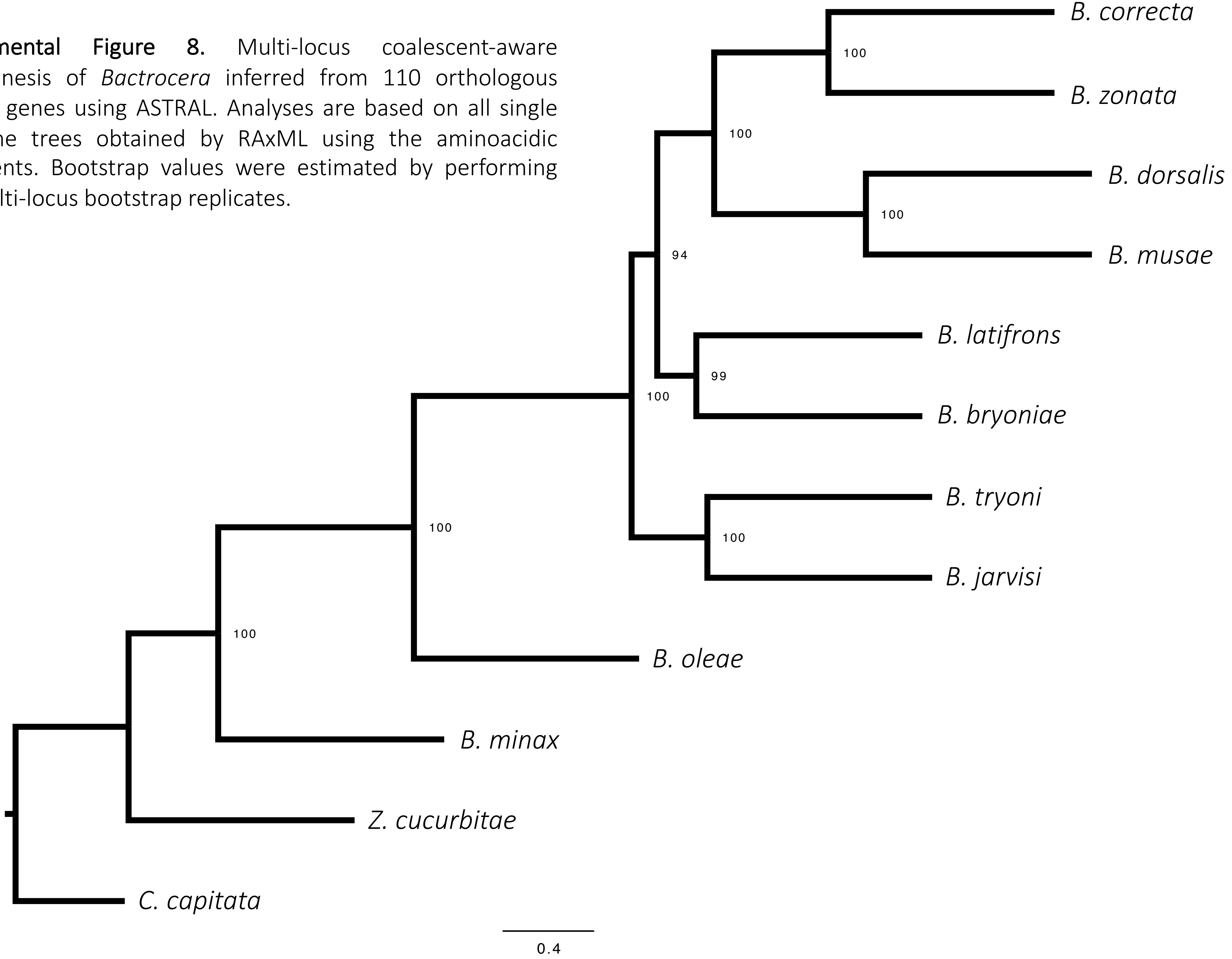

**Supplemental Figure 9.** Multi-locus coalescent-aware phylogenesis of *Bactrocera* inferred from 110 orthologous nuclear genes using ASTRAL. Analyses are based on all single ML gene trees obtained by RAxML using the aminoacidic alignments. Bootstrap values were estimated by performing 100 gene+site resampling.

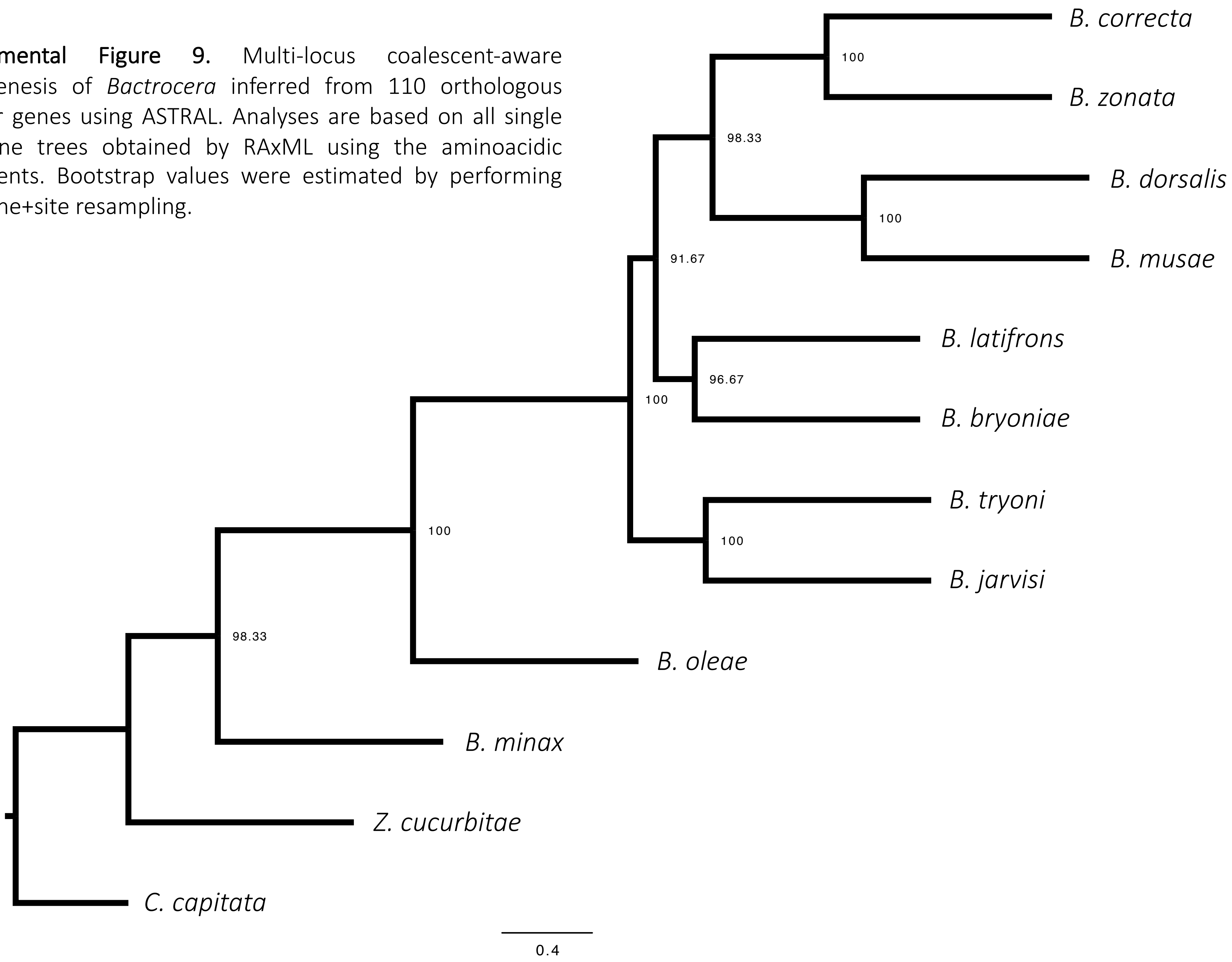

**Supplemental Figure 10.** Multi-locus coalescent-aware phylogenesis of *Bactrocera* inferred from 110 orthologous nuclear genes using ASTRAL. Analyses are based on all single ML gene trees obtained by RAxML using the codon alignments. Bootstrap values were estimated by performing 100 multi-locus bootstrap replicates.

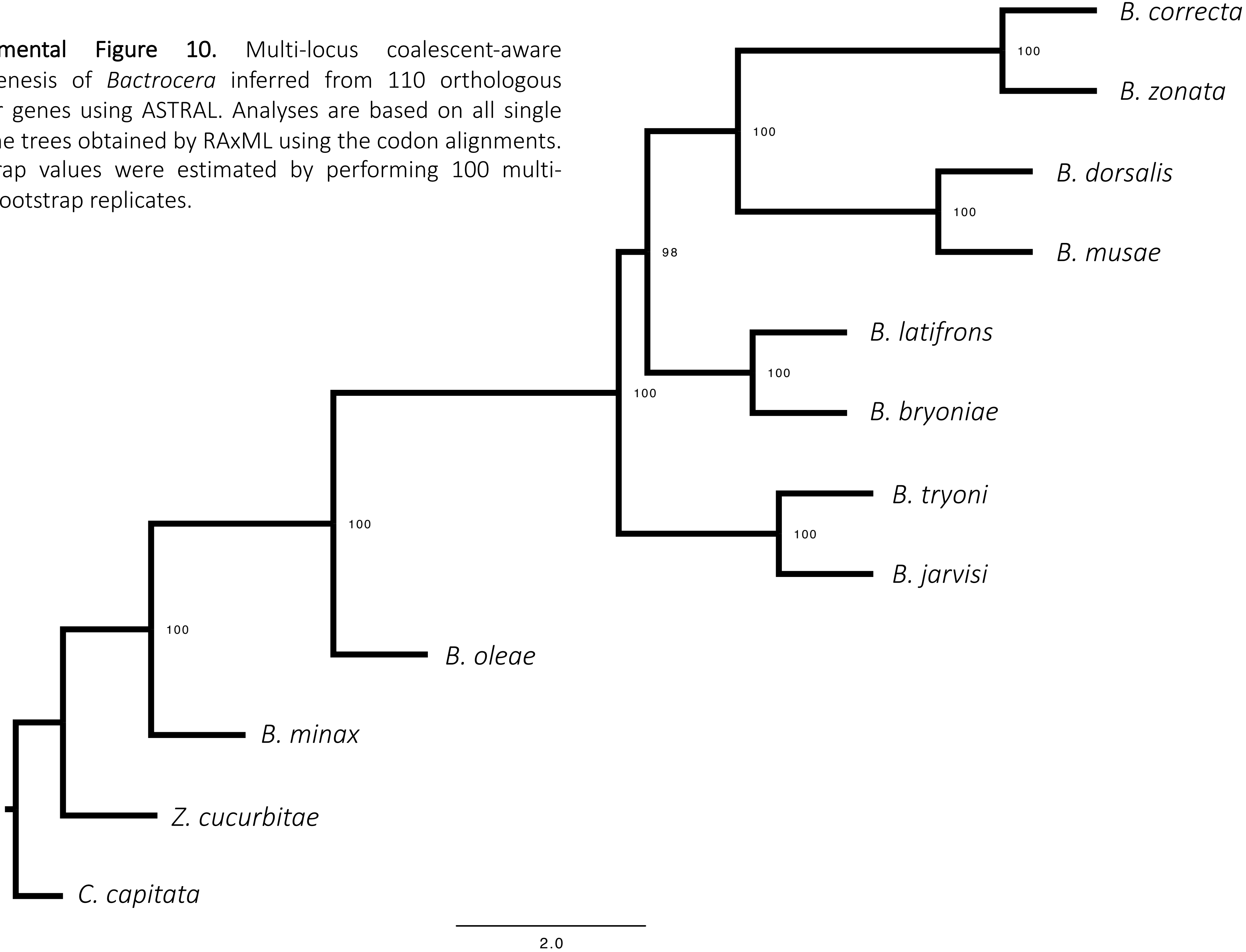

**Supplemental Figure 11.** Multi-locus coalescent-aware phylogenesis of *Bactrocera* inferred from 110 orthologous nuclear genes using ASTRAL. Analyses are based on all single ML gene trees obtained by RAxML using the codon alignments. Bootstrap values were estimated by performing 100 gene+site resampling.

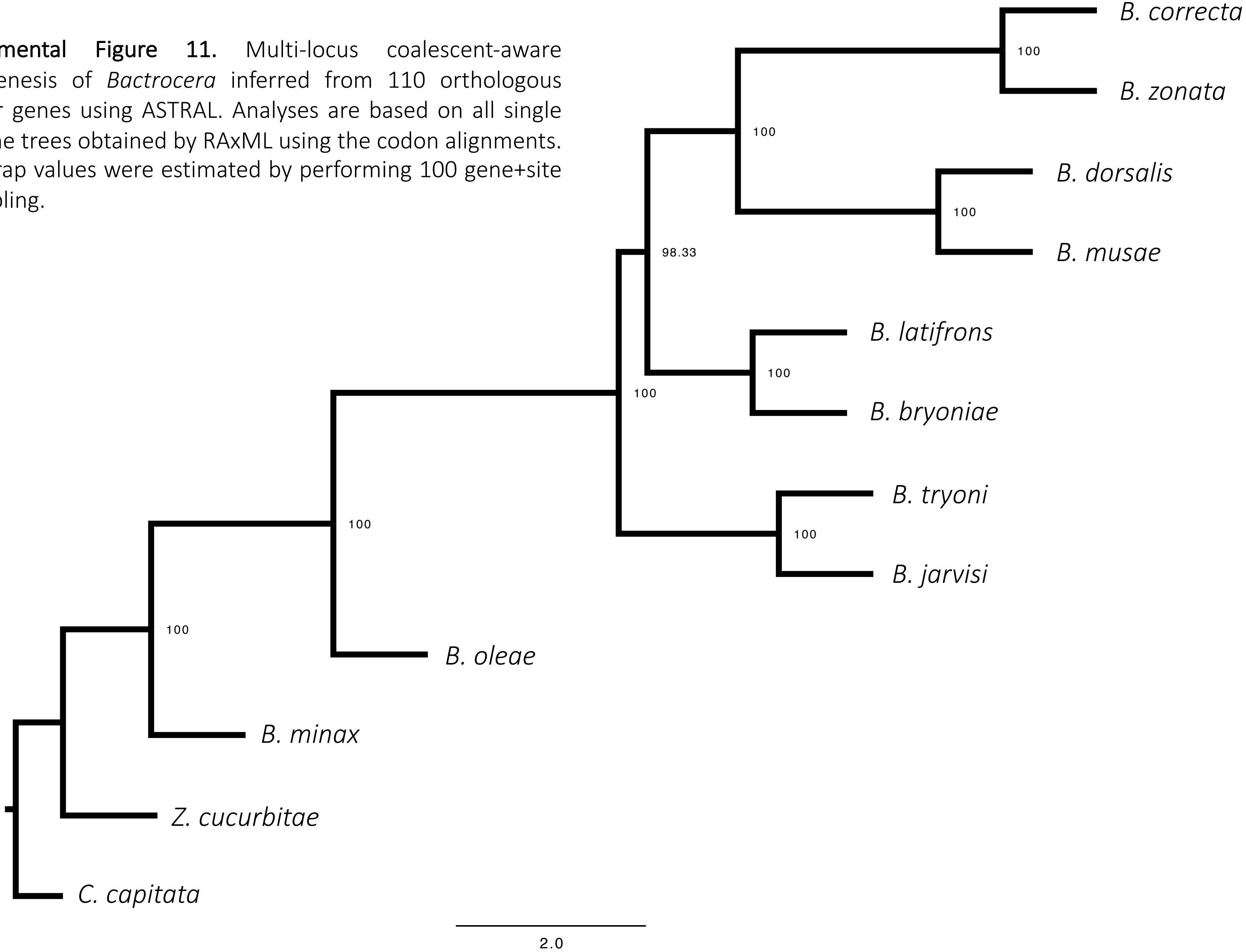

**Supplemental Figure 12.** Multi-locus phylogenesis of *Bactrocera*. Bayesian analyses were obtained by StarBeast2 for each of the 110 amino acidic orthologous genes alignments employing a multispecies coalescent method to estimate the species tree. The site models were linked across the genes.

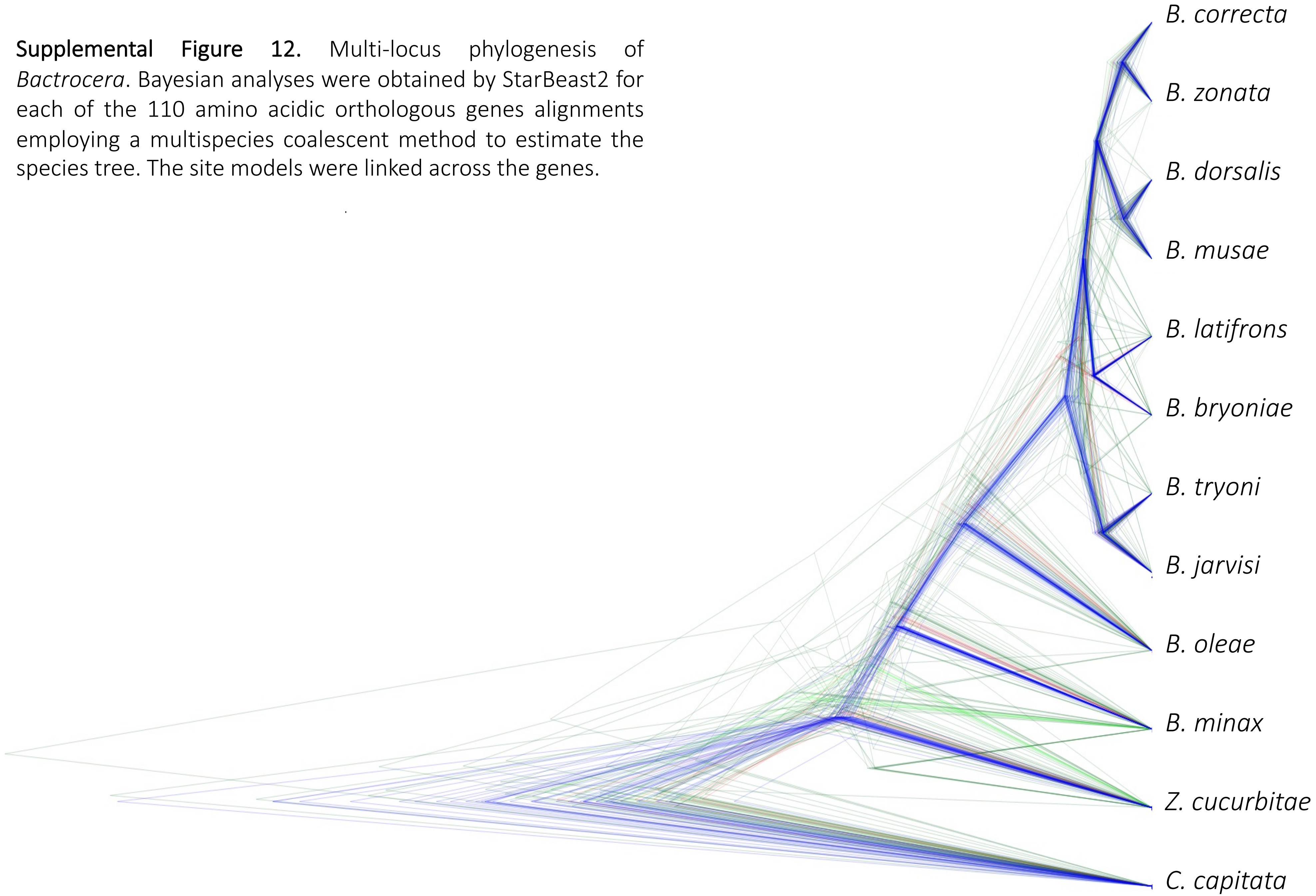

**Supplemental Figure 13.** Multi-locus phylogenesis of *Bactrocera*. Bayesian analyses were obtained by StarBeast2 for each of the 110 codon orthologous genes alignments employing a multispecies coalescent method to estimate the species tree. The site models were linked across the genes.

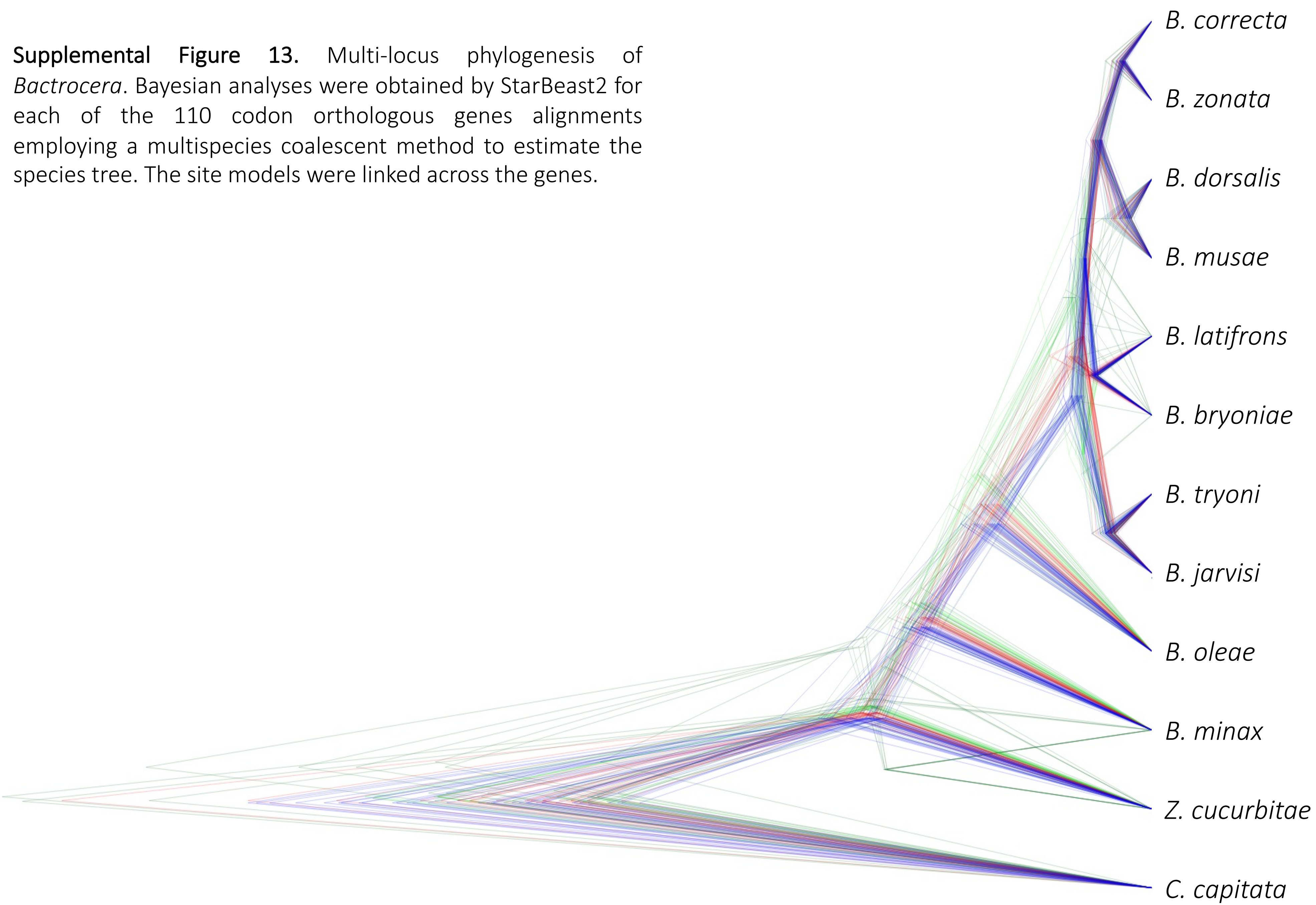

**Supplemental Figure 14.** Molecular time tree of *Bactrocera* obtained by Beast2 using the 4-fold degenerate sites of the concatenated alignments of 110 orthologous nuclear genes (24,885 nt). We used a mutation rate log-normally distributed as prior, a log-normal clock and a Birth-Death model. Mean and 95% confidence interval of the inferred age (corresponding to the blue bars) are reported for each node.

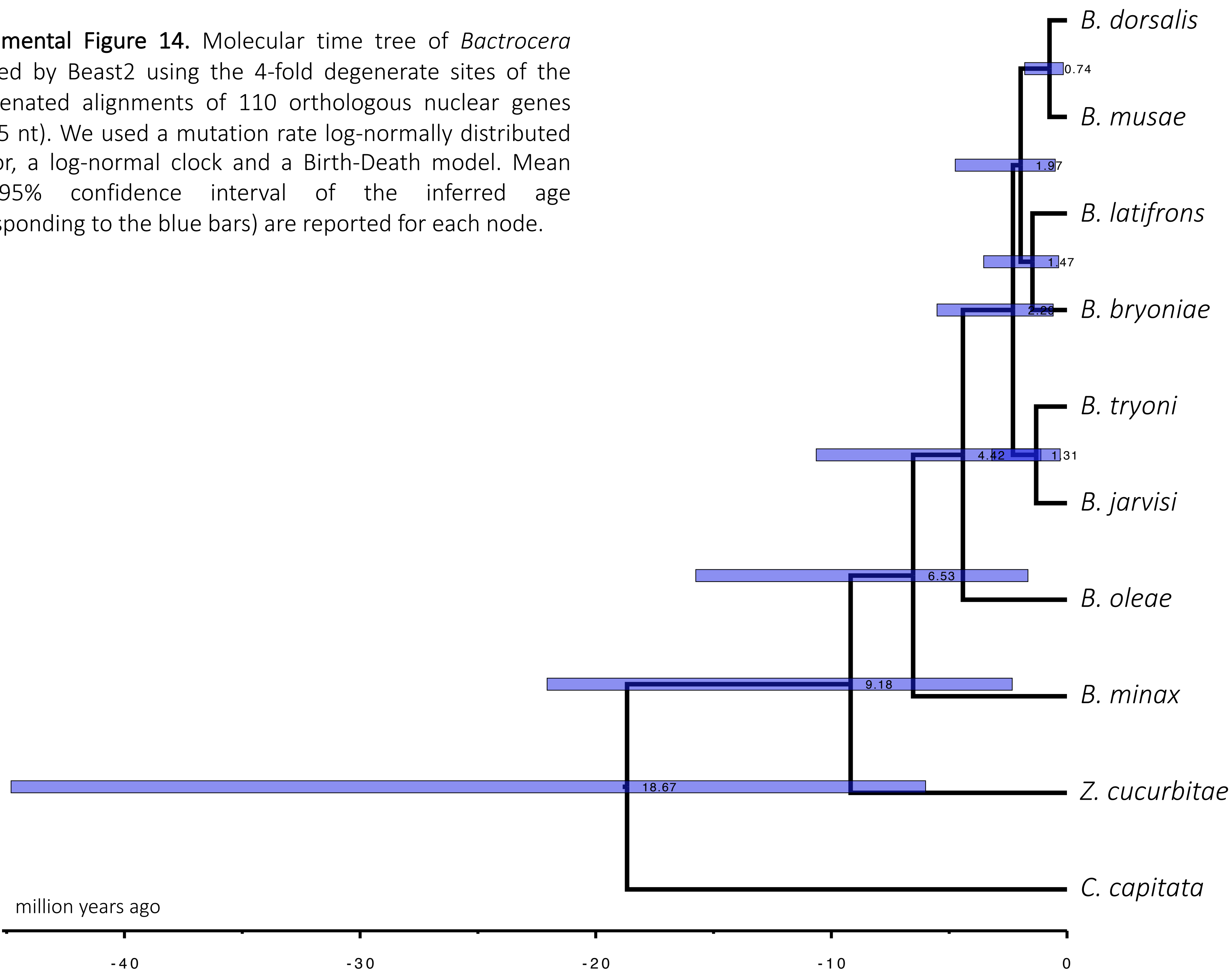

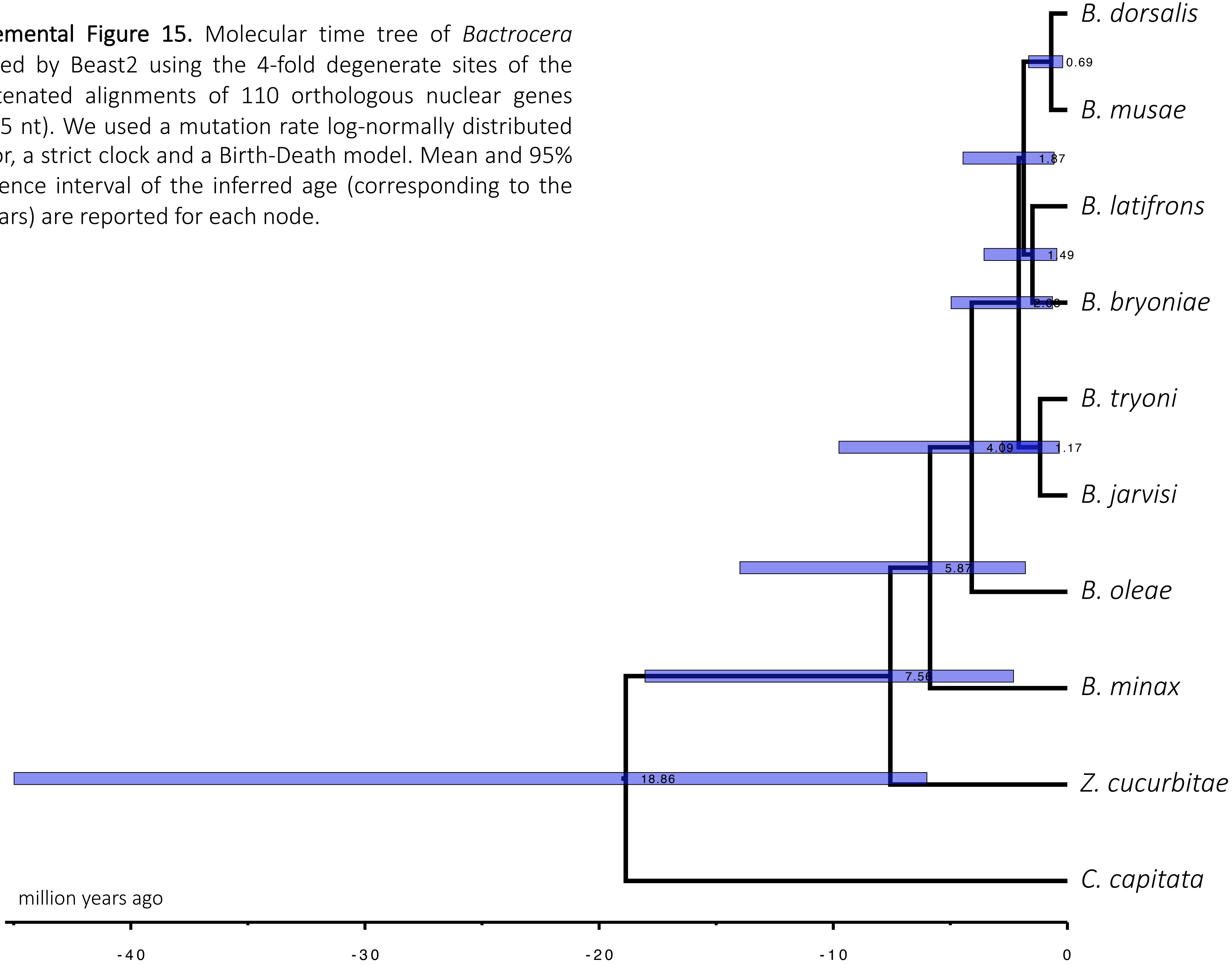


**Supplemental Figure 15.** Molecular time tree of *Bactrocera* obtained by Beast2 using the 4-fold degenerate sites of the concatenated alignments of 110 orthologous nuclear genes (24,885 nt). We used a mutation rate log-normally distributed as prior, a strict clock and a Birth-Death model. Mean and 95% confidence interval of the inferred age (corresponding to the blue bars) are reported for each node.

**Supplemental Figure 16.** Molecular time tree of *Bactrocera* obtained by Beast2 using the 4-fold degenerate sites of the concatenated alignments of 110 orthologous nuclear genes (24,885 nt). We used a mutation rate log-normally distributed as prior, a log-normal clock and a Yule model. Mean and 95% confidence interval of the inferred age (corresponding to the blue bars) are reported for each node.

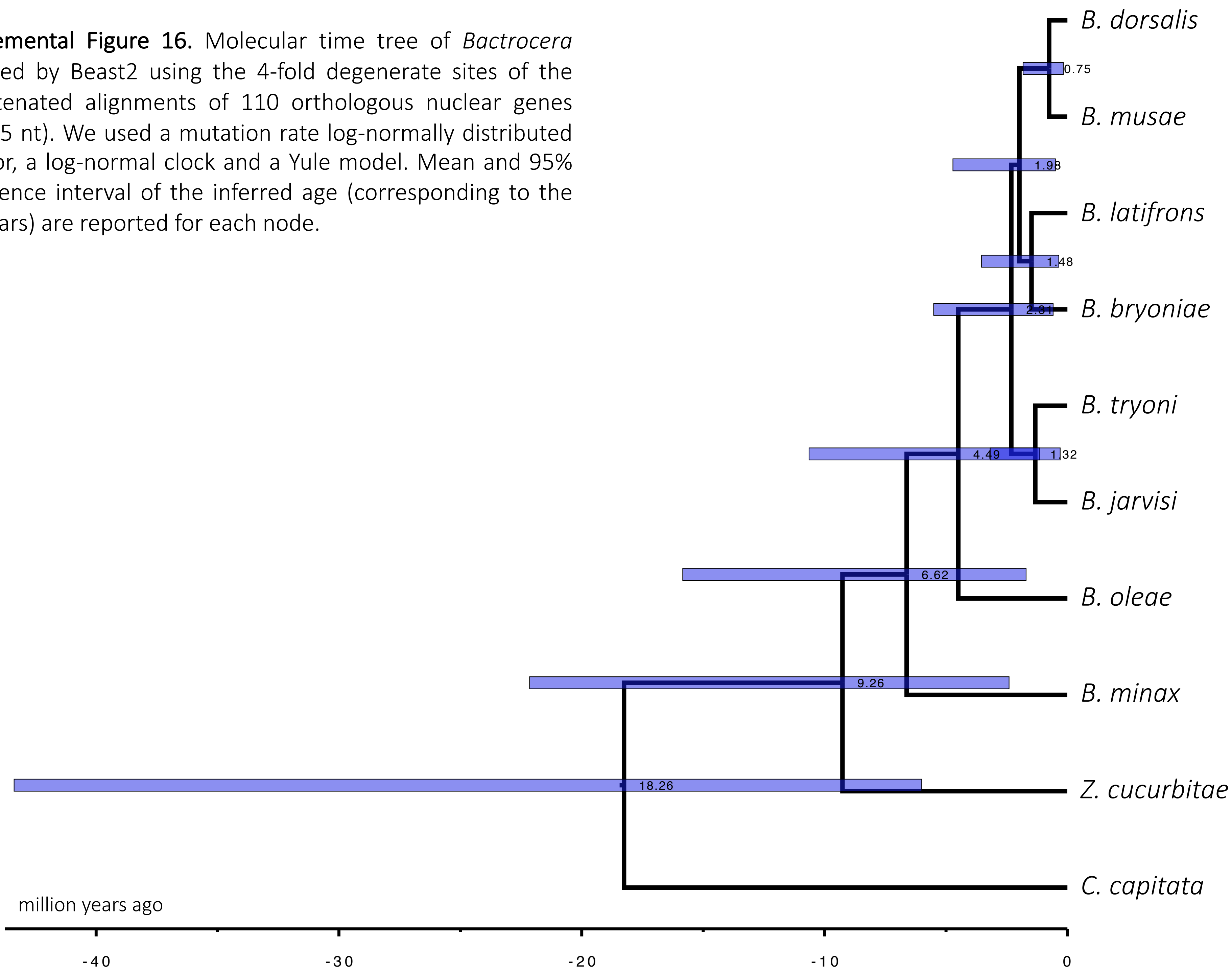

**Supplemental Figure 17.** Molecular time tree of *Bactrocera* obtained by Beast2 using the 4-fold degenerate sites of the concatenated alignments of 110 orthologous nuclear genes (24,885 nt). We used a mutation rate log-normally distributed as prior, a strict clock and a Yule model. Mean and 95% confidence interval of the inferred age (corresponding to the blue bars) are reported for each node.

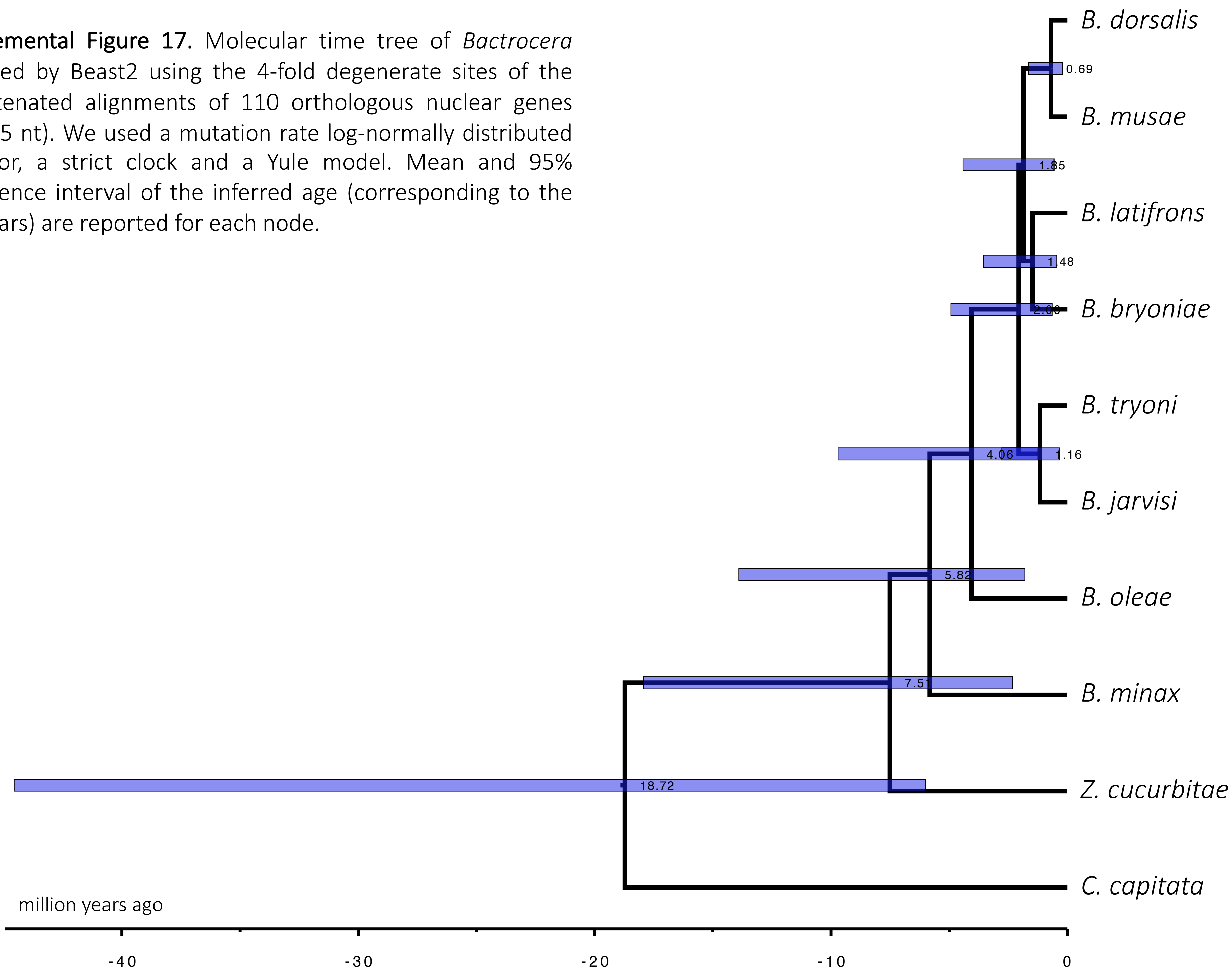

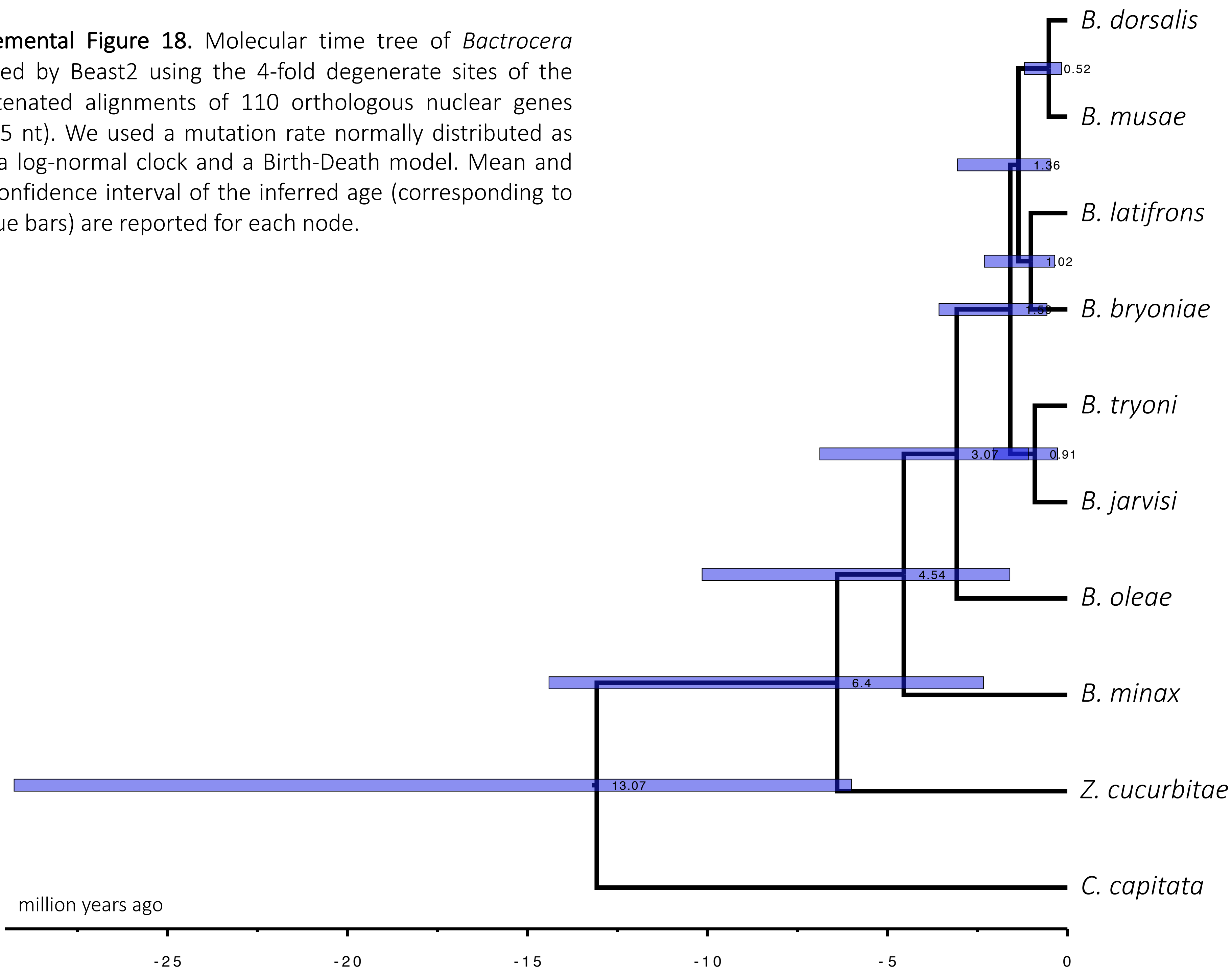


**Supplemental Figure 18.** Molecular time tree of *Bactrocera* obtained by Beast2 using the 4-fold degenerate sites of the concatenated alignments of 110 orthologous nuclear genes (24,885 nt). We used a mutation rate normally distributed as prior, a log-normal clock and a Birth-Death model. Mean and 95% confidence interval of the inferred age (corresponding to the blue bars) are reported for each node.

**Supplemental Figure 19.** Molecular time tree of *Bactrocera* obtained by Beast2 using the 4-fold degenerate sites of the concatenated alignments of 110 orthologous nuclear genes (24,885 nt). We used a mutation rate normally distributed as prior, a strict clock and a Birth-Death model. Mean and 95% confidence interval of the inferred age (corresponding to the blue bars) are reported for each node.

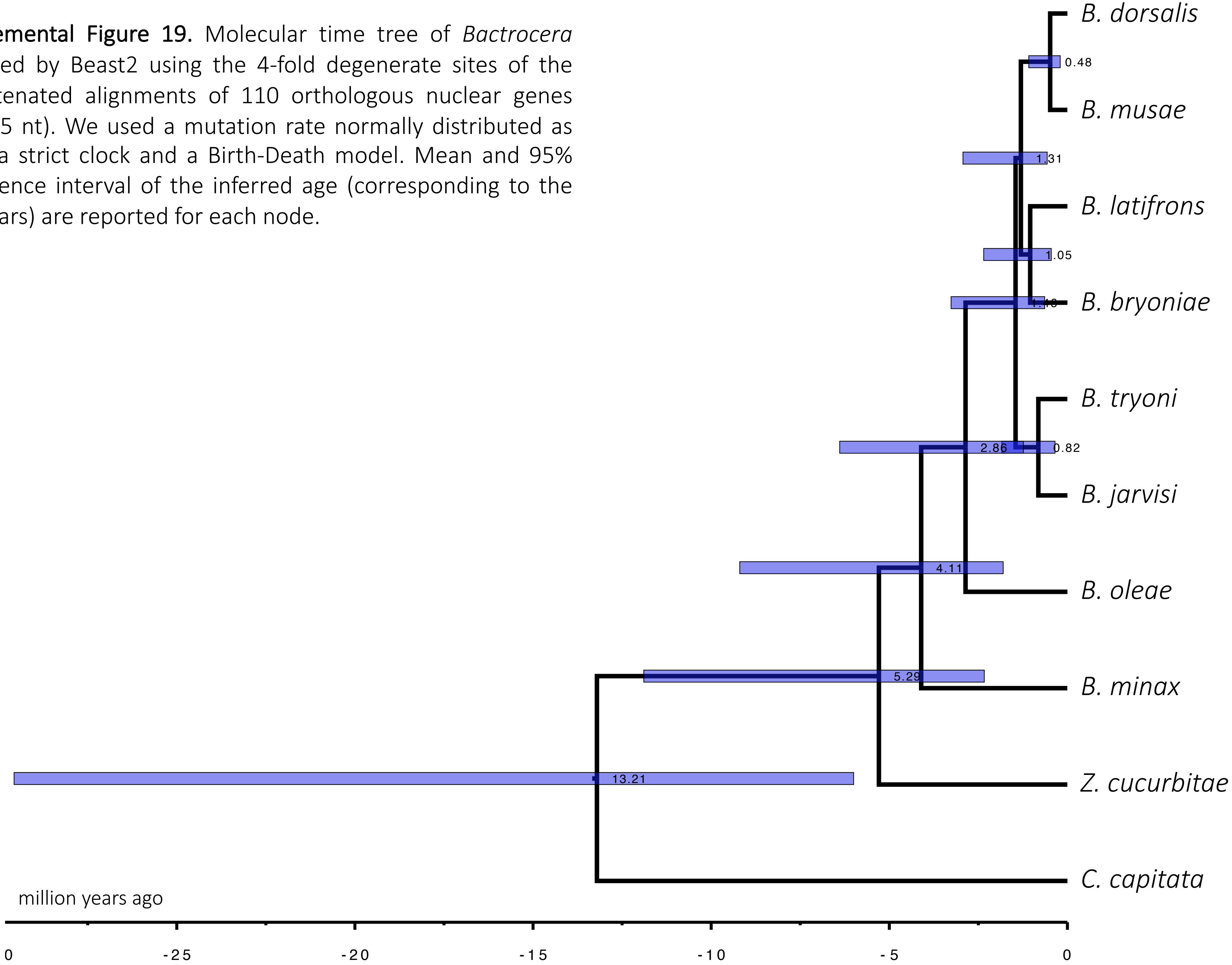

**Supplemental Figure 20.** Molecular time tree of *Bactrocera* obtained by Beast2 using the 4-fold degenerate sites of the concatenated alignments of 110 orthologous nuclear genes (24,885 nt). We used a mutation rate normally distributed as prior, a log-normal clock and a Yule model. Mean and 95% confidence interval of the inferred age (corresponding to the blue bars) are reported for each node.

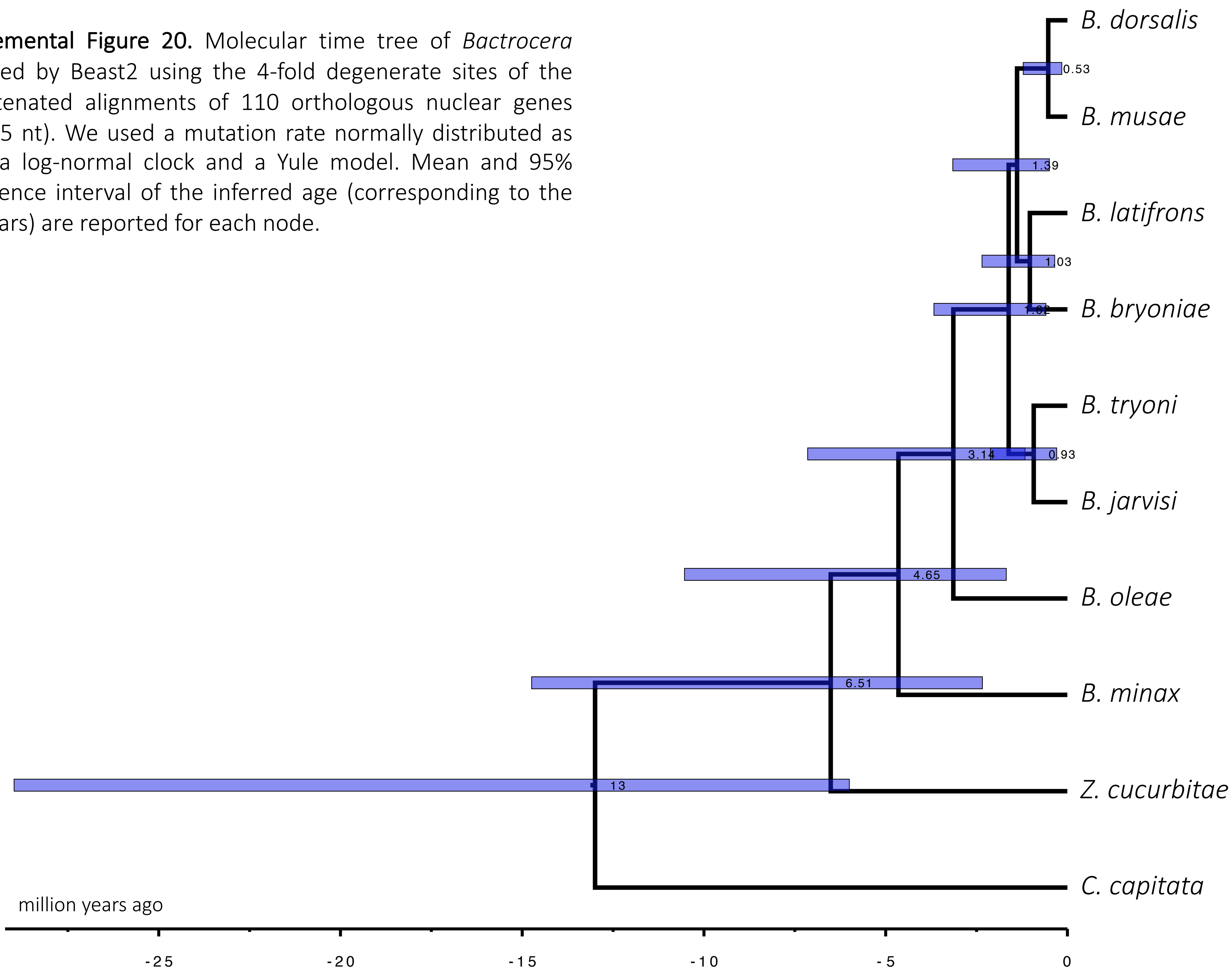

**Supplemental Figure 21.** Molecular time tree of *Bactrocera* obtained by Beast2 using the 4-fold degenerate sites of the concatenated alignments of 110 orthologous nuclear genes (24,885 nt). We used a mutation rate normally distributed as prior, a strict clock and a Yule model. Mean and 95% confidence interval of the inferred age (corresponding to the blue bars) are reported for each node.

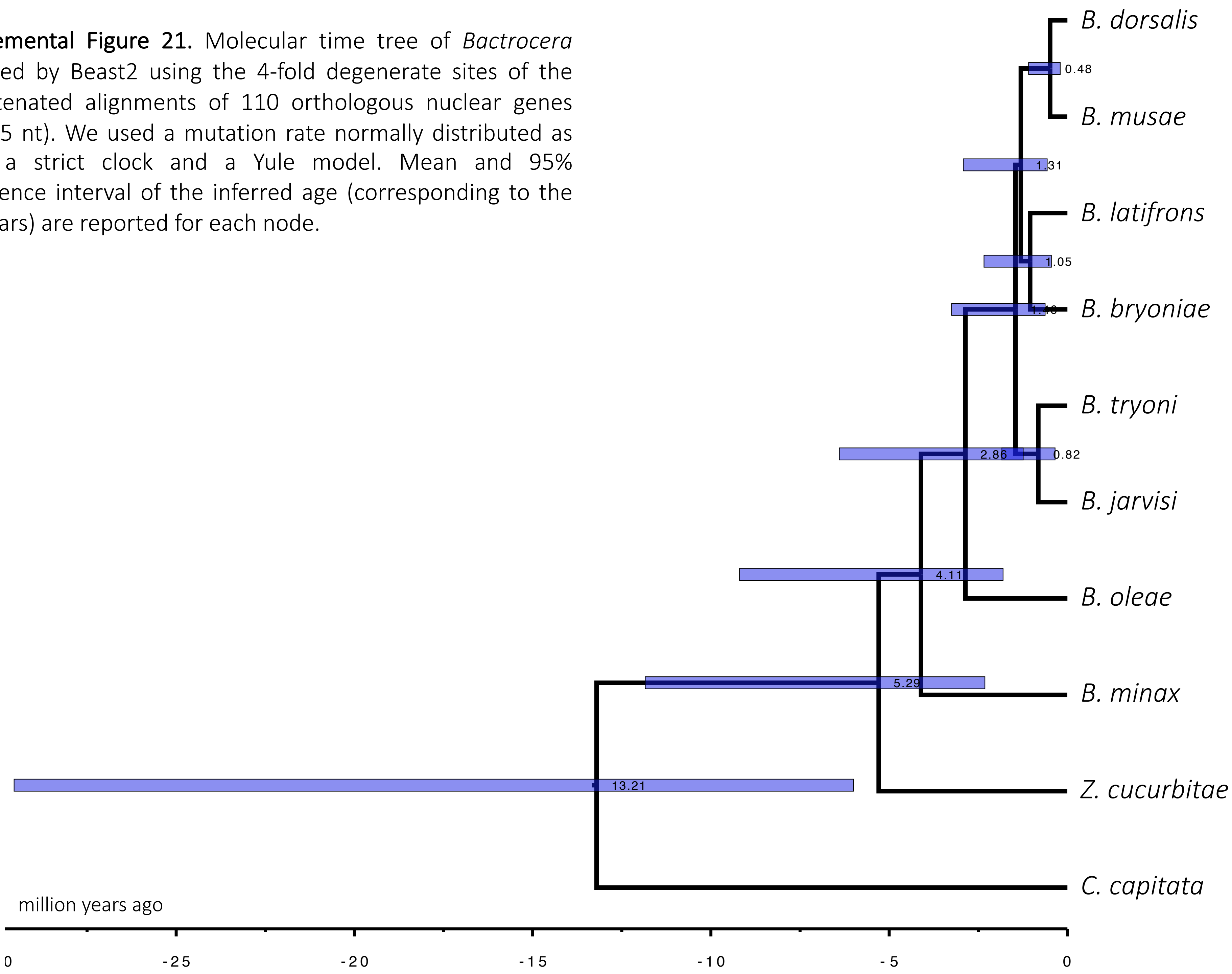
