## Supplemental Table 1 for "The impact of fast radiation on the phylogeny of *Bactrocera* fruit flies"

**Supplementary Table 1. SRA databases used to reconstruct the transcriptome of five *Bactrocera* species.**

| Species | SRA databases (https://www.ncbi.nlm.nih.gov/sra) |
|---|---|
| *Bactrocera bryoniae* | SRX2791703 |
| *Bactrocera correcta* | SRX2013592, SRX2013591, SRX2013590, SRX2013589, SRX2372821, SRX2372818, SRX2372817 |
| *Bactrocera jarvisi* | SRX2791705, SRX697442, SRX697441, SRX697440, SRX697437, SRX697435, SRX697434, SRX697431, SRX697428 |
| *Bactrocera musae* | SRX2791704 |
| *Bactrocera zonata* | SRX2016849, SRX2016848, SRX2016847, SRX2016846 |
