## Supplemental Table 2 for "The impact of fast radiation on the phylogeny of *Bactrocera* fruit flies"

**Supplementary Table 2.** Results of the model selection of the different BEAST analyses used to estimate divergence times.

| | | | *Model 2* | | | | | | | |
|---|---|---|---|---|---|---|---|---|---|---|
| | | | *Mut-normal* | | | | *Mut-lognormal* | | | |
| | | | *yule_strict* | *yule_rlxln* | *bd_strict* | *bd_rlxln* | *yule_strict* | *yule_rlxln* | *bd_strict* | *bd_rlxln* |
| *Model 1* | *Mut-normal* | ***yule_strict*** | - | 62.2 | * | 54.3 | 9.2 | 59.6 | # | 57.2 |
| | | ***yule_rlxln*** | # | - | # | # | # | # | # | # |
| | | ***bd_strict*** | # | 58.6 | - | 50.7 | - | 55.9 | # | 53.5 |
| | | ***bd_rlxln*** | # | 7.9 | # | - | # | 5.2 | # | * |
| | *Mut-lognormal* | ***yule_strict*** | # | 53.1 | # | 45.2 | - | 50.4 | # | 48 |
| | | ***yule_rlxln*** | # | * | # | # | # | - | # | # |
| | | ***bd_strict*** | **6.1** | **68.3** | **9.8** | **60.4** | **15.3** | **65.7** | - | **63.3** |
| | | ***bd_rlxln*** | # | * | # | # | # | * | # | - |

Bayes Factors (BF) were estimated comparing eight BEAST models that combined a mutation prior with either a normal (*Mut-normal*) or lognormal (*Mut-lognormal*) distribution, either a Yule (*yule*) or a Birth-Death (*bd*) model, and either a *strict* or a LOGN relaxed (*rlxln*) clock. Models were compared in a pairwise fashion, by first estimating their Marginal Likelihoods (mL) and corresponding Standard Deviation (SD) and then calculating the Bayes Factors as BF = $mL_1$-$mL_2$, where model 1 and model 2 are those given in the respective row and column. We only report BF values that satisfied the conditions BF > 0 and BF-($SD_1$+$SD_2$) > 0, where $SD_1$ and $SD_2$ are the SD values estimated for model 1 and 2. Highlighted on bold are the values for the model (*Mut-lognormal + bd + strict*) favored over all other seven models. # = BF < 0; * = BF-($SD_1$+$SD_2$) < 0.
